# Resting-state functional dynamics alterations relate to plasma amyloid markers and explain memory impairments in the TgF344-AD model of Alzheimer’s disease

**DOI:** 10.1101/2025.06.01.657264

**Authors:** Alya Al-Awlaqi, Lori Berckmans, Sam De Waegenaere, Sarah Vanherle, Joseph Griffis, Ilse Dewachter, Marleen Verhoye, Mohit H. Adhikari

## Abstract

Resting-state (RS) fMRI studies of Alzheimer’s disease’s (AD) impact on brain function commonly use functional connectivity (FC), ignoring short-timescale network dynamics, captured by co-activation patterns (CAPs), shown to accurately classify transgenic rodents from the wild-type (WT). We acquired high temporal resolution RS-fMRI data in the TgF344-AD rat model at pre-plaque and plaque stages and delineated brain functional alterations using FC and CAPs. We also assessed plaque-stage blood amyloid levels and memory performance in the same animals and investigated the statistical relationship between pathological, RS-functional, and behavioral phenotypes. TgF344-AD (TG) rats had elevated blood amyloid levels, committed more working and reference memory errors and showed reduced hippocampal FC with the lateral cortical and default-mode-like network (DMLN) compared to WT at the plaque stage. They showed DMLN and hippocampal hyper- and hypo-activation at pre- and plaque stages respectively in multiple CAPs. While blood amyloid levels were explained better by plaque-stage, than pre-plaque stage, FC values and CAP activations, it was the pre-plaque stage, more than the plaque stage, CAP activations that accurately explained memory impairments. Our findings not only identify early signatures of AD in brain functional dynamics in this translational rat model but demonstrate their relevance for prognosis of memory deficits.

## Introduction

Alzheimer’s disease (AD) is a neurodegenerative disorder characterized by the accumulation of extracellular amyloid beta (Aβ) plaques and intracellular aggregates of hyperphosphorylated tau proteins (pTau) known as neurofibrillary tangles (NFT). Currently, cerebrospinal fluid (CSF) biomarkers and imaging techniques, especially amyloid-positron emission tomography (PET) and tau-PET, are considered the gold standards for detecting the pathophysiological hallmarks of AD, including Aβ, pTau, and neurodegeneration^1^. Over the past decade, blood-based markers (BBMs) of AD pathology (Aβ38, Aβ40, Aβ42, pTau) as well as of neurodegeneration (neurofilament light chain (NfL) and total tau (tTau)) have gained focus^2^ due to their strong correlation with the CSF measures^3^.

Clinically, AD presents with an insidious onset of episodic memory problems that, due to cortical atrophy, progress to cognitive impairment sufficient to produce functional disability^4^. However, prior to symptoms detectable by cognitive assessments, a prolonged prodromal stage characterized by mild cognitive impairment (MCI) occurs. During this stage, the accumulation of protein aggregates in the brain interferes with neuronal function, eventually leading to neuronal loss^5^. These alterations in neuronal function, particularly at the synapses, result in whole-brain, network-level functional connectivity (FC) changes that can be measured with resting-state functional magnetic resonance imaging (RS-fMRI).

RS-fMRI has been instrumental in providing a network perspective of the functional impact of several neurological disorders such as stroke^6^, autism spectrum disorders^7^, and neurodegenerative diseases^8,9^. In the context of AD, increased Aβ levels have been shown to correlate with alterations in FC within the default mode network (DMN), the most prominent resting-state network (RSN)^10^. We have found, in the TgF344-AD rat model, a reduced RS-FC limited to a few specific connections at 6-months and becoming more widespread at 10-months^11^. Beyond traditional FC, temporal analysis of RS-activity has revealed transient, recurring brain states known as co-activation patterns (CAPs), which reflect the dynamic organization of RSNs ^7,12^. Our previous findings in the Tg2576 AD mouse model demonstrated that CAPs effectively distinguish 18-month-old AD mice from WT controls^13^. Additionally, in a longitudinal study of a Huntington’s disease (HD) mouse model, we found that the spatial properties of CAPs not only reveal very early alterations in the HD model mice, but they can also predict HD mice from WT more accurately than FC, and capture phenotypic progression^14^. Most recently, we utilized the single time-frame resolution of CAPs to examine the inter-CAP transition probability changes in the TgF344-AD rat model of AD at pre- and early-plaque stages^15^. The TG rats showed differential transition probabilities within the same CAP-pair at both stages, whereas the WT rats did so only at the pre-plaque stage. Thus, these sensitive, analytic measures based on RS-fMRI can reveal nuanced and early changes in brain functional dynamics due to AD. However, it is unclear how these measures are related to markers of AD pathology and/or alterations in memory/cognitive performance, and whether the functional changes particularly at early stages have any prognostic relevance.

In this study, we aimed to investigate the relationship between BBMs, memory alterations and brain’s functional dynamics measured in the same cohort of TgF344-AD and WT rats. We assessed the memory performance and obtained BBMs in these animals at 10.5-11 months of age, which corresponds to the plaque stage in this model. We acquired RS-fMRI data at 4 (pre-plaque) and 10 months of age and used FC and CAPs to characterize changes in the brain’s functional architecture and their progression with age. We hypothesized that both the BBMs and memory impairments in the model rats could be explained by RS-fMRI measures, particularly by RS-CAPs at the pre-plaque stage, demonstrating their prognostic value.

## Results

### Pathological and behavioral phenotypes at the plaque stage in the TG animals

We assessed AD pathology in the TG animals by obtaining blood-plasma levels of amyloid markers (Aβ38, Aβ40, and Aβ42) at 11 months (M) of age and comparing them to those from the WT animals. As expected, the levels for all three markers were significantly higher in the TG animals (p < 0.01 for Aβ38, unpaired T-test, p < 0.0001 for Aβ40 and Aβ42, Mann-Whitney U test, Figure 1). We also assessed the levels of NfL and GFAP in the blood samples of these animals. While levels of NfL were not significantly different between the two genotype groups, the values of GFAP for all animals were below the detection range.

**Figure 1:**
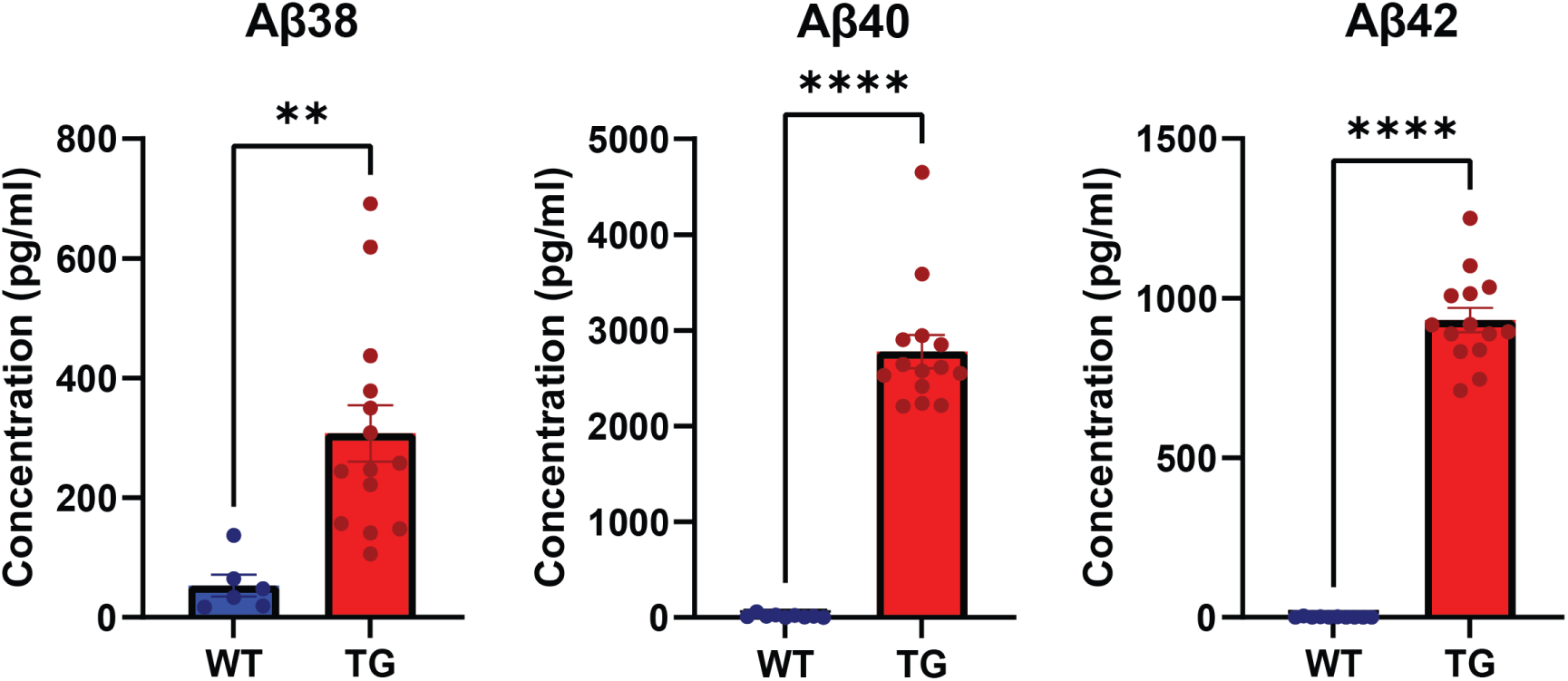
Amyloid marker values in the blood-plasma samples of 11M old WT and TG animals. **: p < 0.01; unpaired T-test, ****: p < 0.0001: Mann-Whitney U test.

We analyzed and identified the changes in the performance on memory tasks in the TG animals compared to the WT at 10.5M. Of the 10 measures we obtained over the entire acquisition period of 10 days, TG animals took longer to complete the task (Figure 2A, p = 0.0028, main genotype effect), spent more time and travelled a higher percentage of distance in the non-baited (wrong) arms (Figure 2H, 2I, p = 0.0050, main genotype effect), and made a higher number of reference memory errors (Figure 2B, p = 0.0271, main genotype effect) compared to WT rats. In the last 3-4 days of acquisition, compared to the first acquisition day, WT animals made fewer working memory errors (Figure 2C, day 10 vs day 1 p = 0.0006, FDR corrected), entered fewer arms (Figure 2D, day 10 vs day 1 p = 0.0448, FDR corrected) and travelled a shorter distance (Figure 2E, day 10 vs day 1 p = 0.0097, FDR corrected). In contrast, TG animals did not improve their performance over time and hence in the last 3-4 days, showed a significantly higher percentage of working memory errors (Figure 2C, day 10 p = 0.0006, FDR corrected), entered more arms (Figure 2D, day 10 p = 0.0020, FDR corrected), and travelled longer total distances (Figure 2E, day 10 p = 0.0003, FDR corrected) compared to the WT rats. Mean speed showed a significant interaction effect and subsequent post hoc comparisons showed a significant time effect for both the WT and TG animals (Figure 2F, day 10 vs day 1 p < 0.0001 (WT), p < 0.0001 (TG), post hoc, FDR corrected) while the latency to enter the first non-baited arm was significantly shorter in the TG on day 10 compared to the WT (Figure 2G, p = 0.0008, FDR corrected).

**Figure 2:**
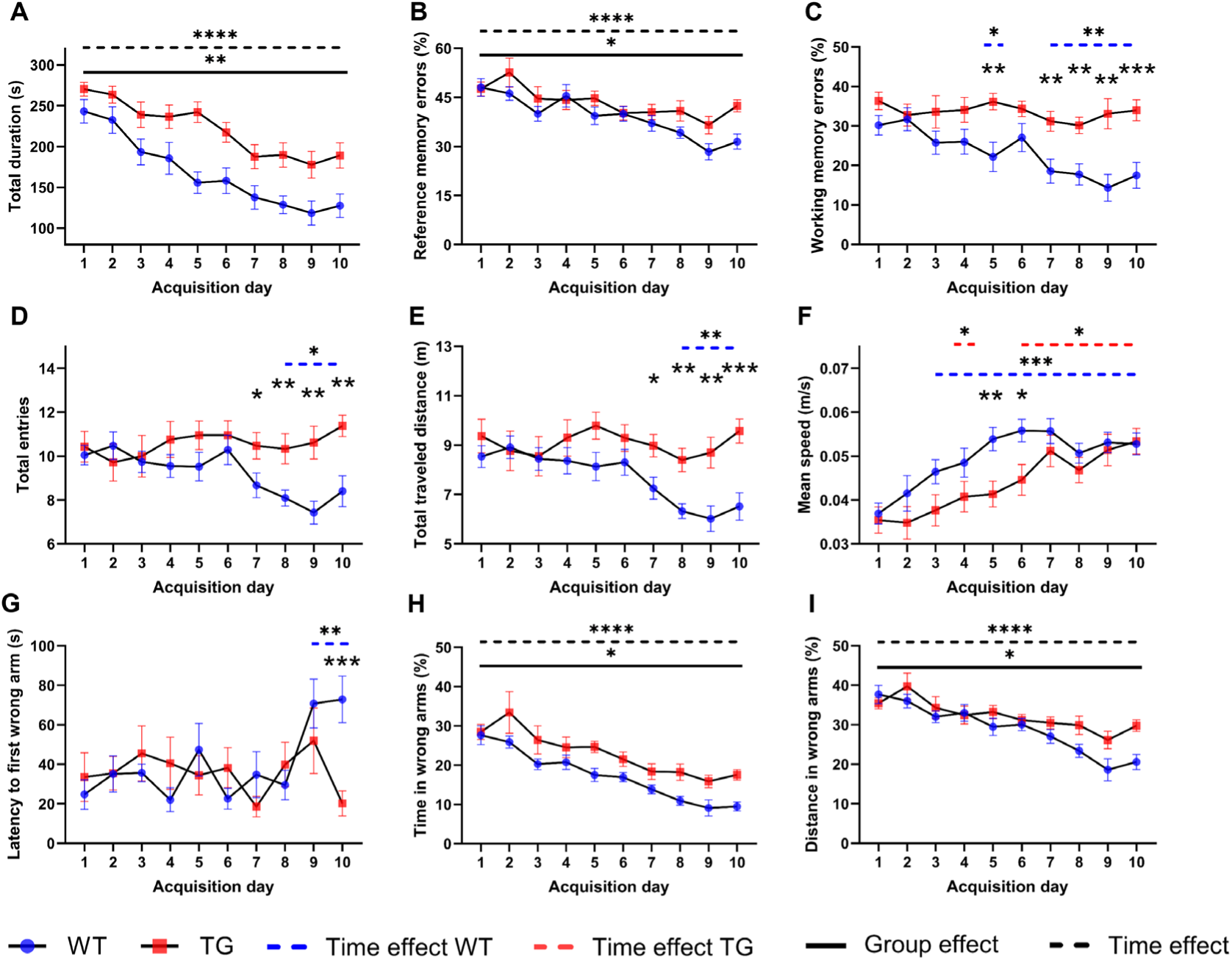
Comparison of nine memory task performance measures – total duration (A), reference memory errors (B), working memory errors (C), total entries (D), total travelled distance (E), mean speed (F), latency to enter the first non-baited (wrong) arm (G), time spent (H) and distance travelled (I) in the non-baited arms by 10.5M old WT and TG animals in radial arm maze over 10 days. *: p < 0.05; **: p < 0.01; ***: p < 0.001; post-hoc comparison between the genotypic groups on specified acquisition days following a significant group x time interaction effect. Blue and red dashed lines with asterisks indicate significant performance differences on post-hoc comparisons between the specified days (extent of the line) and day 1, following a significant genotype x time interaction effect, in the WT and TG animals respectively. Black solid line: significant main group effect; Black dashed line: significant main time effect on the two-way ANOVA.

### Functional changes at the pre-plaque and plaque stages in the TG animals

Next, we analyzed the RS-fMRI data acquired in these animals at 4M and 10M using both the traditional static FC approach as well as the more advanced dynamic FC approach of CAPs. Figures 3A-B display the mean ROI-based FC matrices of both genotypic groups at 4M and 10M, respectively. Visual inspection indicates high FC within the DMLN, hippocampal network (Hipp), sensory (Sens) network, lateral cortical network (LCN) and high anticorrelation between the LCN and the DMLN, Hipp and Sens networks, especially in the WT group at 10M. FC of several ROI-pairs showed a significant interaction effect, but post hoc comparisons showed no significant inter-genotypic differences at either 4M or 10M (p > 0.05, FDR corrected; Supplementary Figure 1A). Functional correlation of CA1 and Ent of Hipp network with Rsp and TeA, respectively, of the DMLN in the right hemisphere and anticorrelation between several regions of the LCN and Hipp network was increased at 10M compared to 4M only in the WT group while no inter-age differences were found to be significant in the TG group (Supplementary Figure 1B). We found a significant age effect in almost all ROI-pairs without a significant interaction effect and a significant group effect in four pairs. However, neither the difference between ages nor the inter-group difference remained significant after correcting for post-hoc comparisons (p > 0.05; FDR corrected; Supplementary Figure 1C). Figures 3C-F display the corresponding mean network-based FC matrices in WT and TG groups at 4M and 10M. We found a significantly lower correlation and anticorrelation of the hippocampal network with the DMLN and the LCN (Figure 3G), respectively, in the TG group at 10M, compared to WT (p = 0.015, for both Hipp-DMLN and Hipp-LCN; FDR corrected for 2 comparisons), and in the 4M WT group compared to the 10M WT group (Figure 3H, p =7E-4 (Hipp-DMLN), p = 1E-3 (Hipp-LCN); FDR corrected for 2 comparisons). All within-network FC, except the sub-cortical network, showed a significant increase at 10M compared to 4M irrespective of groups (Figure 3I, p < 0.05, FDR corrected). The magnitude of 6 between-network FCs was also found to be higher at 10M compared to at 4M (Figure 3I, p < 0.05, FDR corrected).

**Figure 3:**
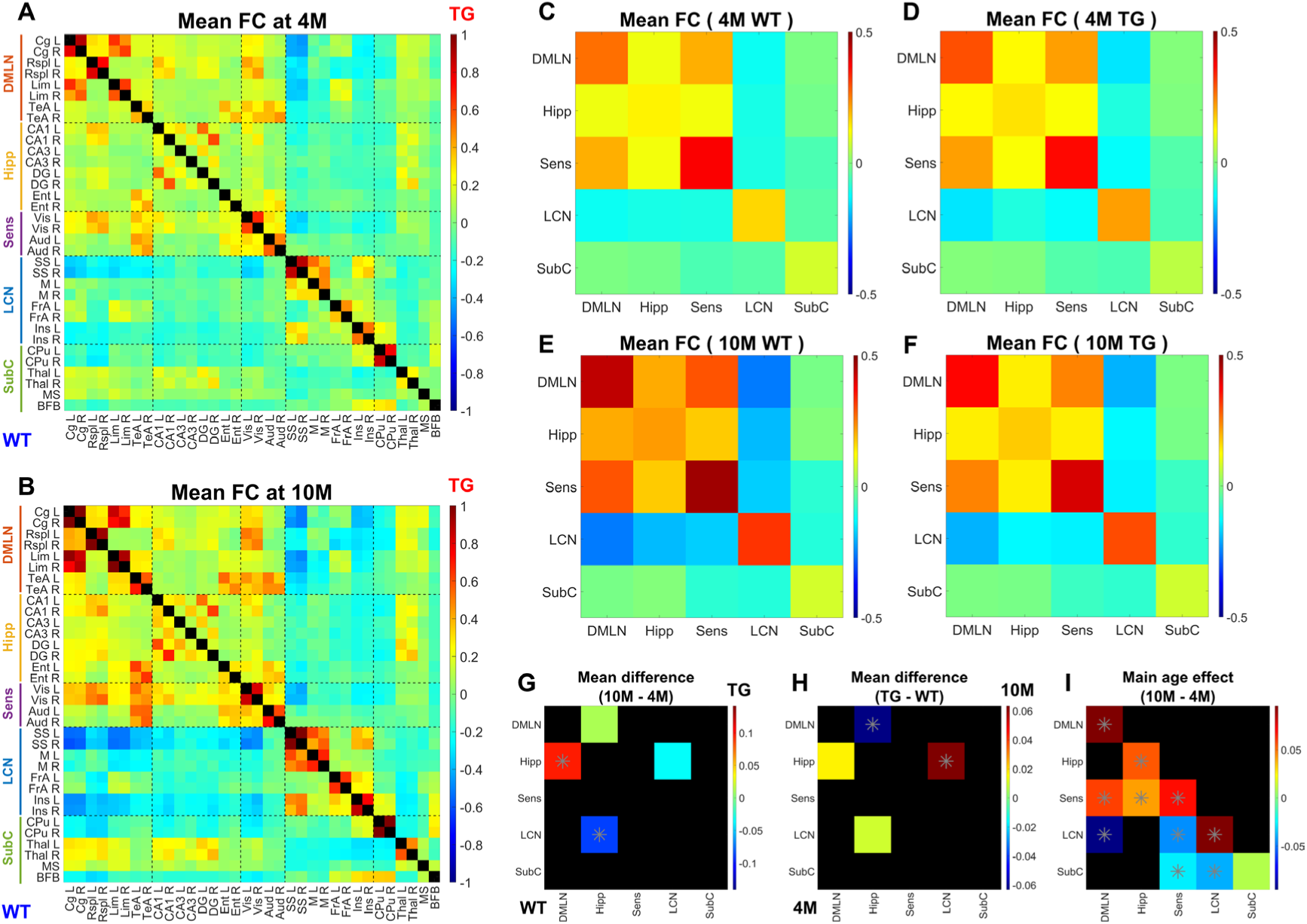
A, B: Mean ROI-based FC between 34 ROIs in the WT (below the diagonal) and in the TG (above the diagonal) at 4M (A) and 10M (B). **C-F:** Mean network-level FC for five RSNs in 4M WT (C), 4M TG (D), 10M WT (E), and 10M TG (F) groups. **G, H, I:** Results of two-way repeated measures ANOVA analysis on the network-level FC values. G & H show the inter-genotype difference (G; 4M/10M: below/above the diagonal) and inter-age difference (H; WT/TG: below/above the diagonal) for the Hipp-LCN FC and Hipp-DMLN FC that showed a significant interaction effect. Asterisk: p < 0.05, FDR corrected for 4 post-hoc comparisons. I: the inter-age difference for all inter-and intra-network FC values that showed a significant age effect. Asterisk: p < 0.05, FDR corrected for 12 post-hoc comparisons.

Next, we applied CAPs analysis to examine the functional changes in instantaneous brain states in the TG animals compared to the WT controls. After concatenating the image series from all subjects at both ages and clustering them across a range of 2 to 20 clusters, we determined that six CAPs optimally accounted for the variance in the image series. (Supplementary Figure 2A-B). Figures 4 A, D, G, J, M, and P show color-coded one-sample T-test maps for significantly activated/deactivated voxels in these six CAPs (p < 0.01, Bonferroni corrected). The DMLN CAP (Figure 4A) is characterized by the co-activation of the DMLN (cingulate, retro splenial cortices), sensory (auditory, visual cortices) regions, and basal forebrain with simultaneous co-deactivation of the LCN (motor, somatosensory and insular cortices) regions. The LCN CAP (Figure 4D) presents an anticorrelated pattern to the DMLN CAP (Supplementary Figure 2C). The next two CAPs show fragmented LCN activations with simultaneous DMLN deactivation. The Rostral LCN CAP (Figure 4G) features co-activation of the rostral and lateral parts of LCN regions along with the CPu while the Caudal LCN CAP (Figure 4J) shows co-activation of caudal & medial LCN regions thus an anticorrelated pattern to the Rostral LCN CAP (Supplementary Figure 2C). The CPu CAP (Figure 4M) is characterized by co-activation of the CPu and the thalamus and co-deactivation of particularly the rostral LCN regions. The DMLN Hipp CAP (Figure 4P) shows co-activations of the DMLN, hippocampus and the sensory regions with simultaneous co-deactivated CPu and LCN regions. The CPu and the DMLN Hipp CAPs did not show high anticorrelation (Supplementary Figure 2C).

**Figure 4:**
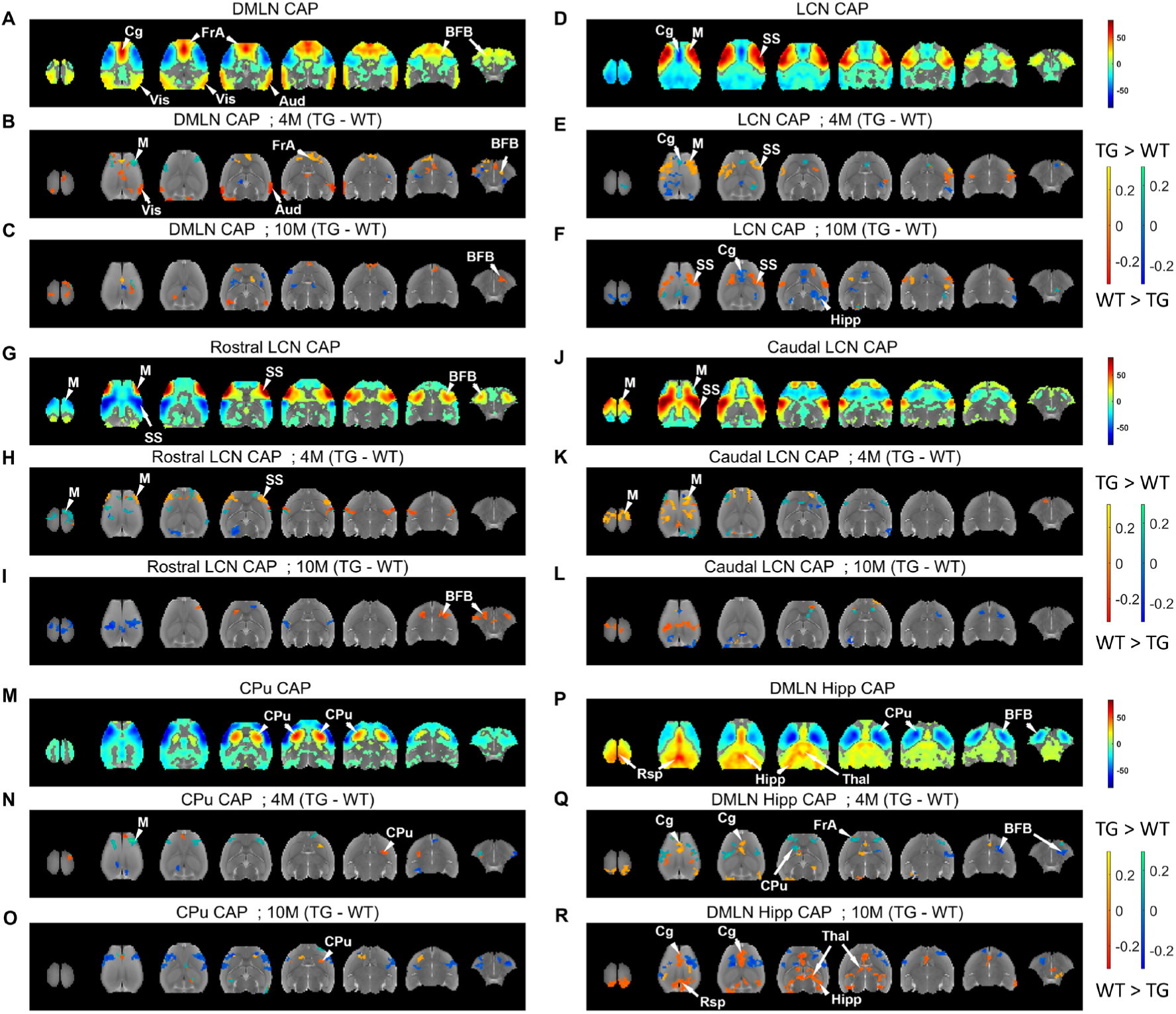
Voxel wise (de)activations of combined CAPs and comparisons of mean (de)activations between WT and TG at 4M and 10M. **A, D, G, J, M, P:** Color-coded one-sample T-test statistic values for voxels whose z-scored blood-oxygen-level-dependent (BOLD) signal averaged across timeframes belonging to each CAP is significantly (p<0.01, Bonferroni corrected) higher (yellow to red, activation) and lower (cyan to blue, deactivation) than zero. **B/C, E/F, H/I, K/L, N/O, Q/R:** Genotypic difference in (de)activation in CAPs 1-6 at 4M/10M for voxels showing significance (p < 0.05, FDR corrected) in post-hoc comparisons following a significant interaction effect (p < 0.05, two-way ANOVA). These two-way ANOVA comparisons are performed for voxels that show significant (de)activation in at least one age/genotypic combination for each CAP. In each post-hoc comparison maps, red to yellow and blue to green color bars represent the inter-genotypic differences for voxels that are co-activated and co-deactivated, respectively, in the WT group. Thus, positive (in yellow and green) and negative (in red and blue) differences correspond to a significantly higher and lower magnitude of (de)activation, respectively, in the TG group vis-à-vis the WT group.

We first assessed the genotype, age, and genotype x age interaction effects on the temporal properties of CAPs. The duration of the Rostral and Caudal LCN CAPs was significantly lower at 10M compared to 4M in both genotypic groups. Additionally, the LCN CAP showed significantly reduced duration in the TG group at 10M (p<0.05, FDR corrected, Supplementary Figure 3A). Occurrence percentages for both the Caudal LCN and CPu CAPs were, respectively, significantly lower and higher, at 10M compared to 4M (p<0.05, two-way ANOVA, Supplementary Figure 3B). Next, we examined the genotype, age and genotype x age interaction effects on the voxel-level activations in CAPs. Figures 4B and 4C display the age-specific genotypic difference in the DMLN CAP activation levels for voxels that showed a significant interaction effect (p < 0.05, two-way ANOVA) and a significant post-hoc genotypic difference (p < 0.05, FDR corrected for the number of voxels with significant interaction effect) at 4M and 10M, respectively. Figures 4E and 4F, 4H and 4I, 4K and 4L, 4N and 4O, 4Q and 4R represent similar intergenotypic differences at 4M and 10M for other CAPs respectively. Most prominently, at 4M, we found a common thread across all CAPs. Most of the voxels with significant intergenotypic difference in their (de)activation, showed significant hyper activation (represented in yellow) or hyper deactivation (represented in green) in the TG animals vis-à-vis the WT. At 10M, on the other hand, our prominent finding across all CAPs was a significant hypoactivation (red) and hypo-deactivation (blue) in most voxels in the TG animals. These effects were most robustly observed in the LCN CAP, the Caudal LCN CAP, and the DMLN Hipp CAP. In the LCN CAP, at 4M, we found prominent hyperactivation in the LCN regions and a hyper-deactivation in the Cg (Figure 4E) which reversed to hypoactivation at 10M (Figure 4F). In the Caudal LCN CAP, parts of LCN regions showed hyper- and hypoactivation in the TG animals at 4M and 10M respectively. Similarly, the DMLN Hipp CAP showed hyperactivation in the Cg at 4M (Figure 4Q) while at 10M, it showed hypoactivation in the DMLN regions, parts of Hipp, and hypo deactivation in the LCN regions, (Figure 4R) in the TG animals vis-à-vis the WT. Interestingly, in the DMLN Hipp CAP, the BFB showed hypo deactivation in the TG group at both 4M and 10M (Figure 4Q, 4R).

### Relationship between functional changes and AD pathology

In the next step, we assessed whether the levels of amyloid markers in blood plasma in the TG animals at the plaque stage could be explained, statistically, using functional markers such as region of interest (ROI)- and network-based FC, and spatial activation of CAPs at both pre-plaque and plaque stages. Figure 5 shows the *R*^2^, i.e., the variance in each of the three amyloid markers in the TG animals explained by their ROI-based FC (Figure 5 A-C) and voxel-wise activations in all six CAPs (Figure 5 D-F) at 4M and 10M. Here, we used the FC between all 34 ROIs, considered in Figures 3 A-B and T-statistics for all voxels found to be significant in combined CAPs (colored voxels in Figures 4 A, D, G, J, M, P). The *R*^2^ of the fitted statistical models using both ROI-level FC and CAPs for each amyloid marker always exceeded 0.5 and was higher at 10M with narrower confidence intervals compared to 4M (Figure 5 A-F). We found low *R*^2^ (< 0.3) values when network-based FC values were used (data not shown).

**Figure 5:**
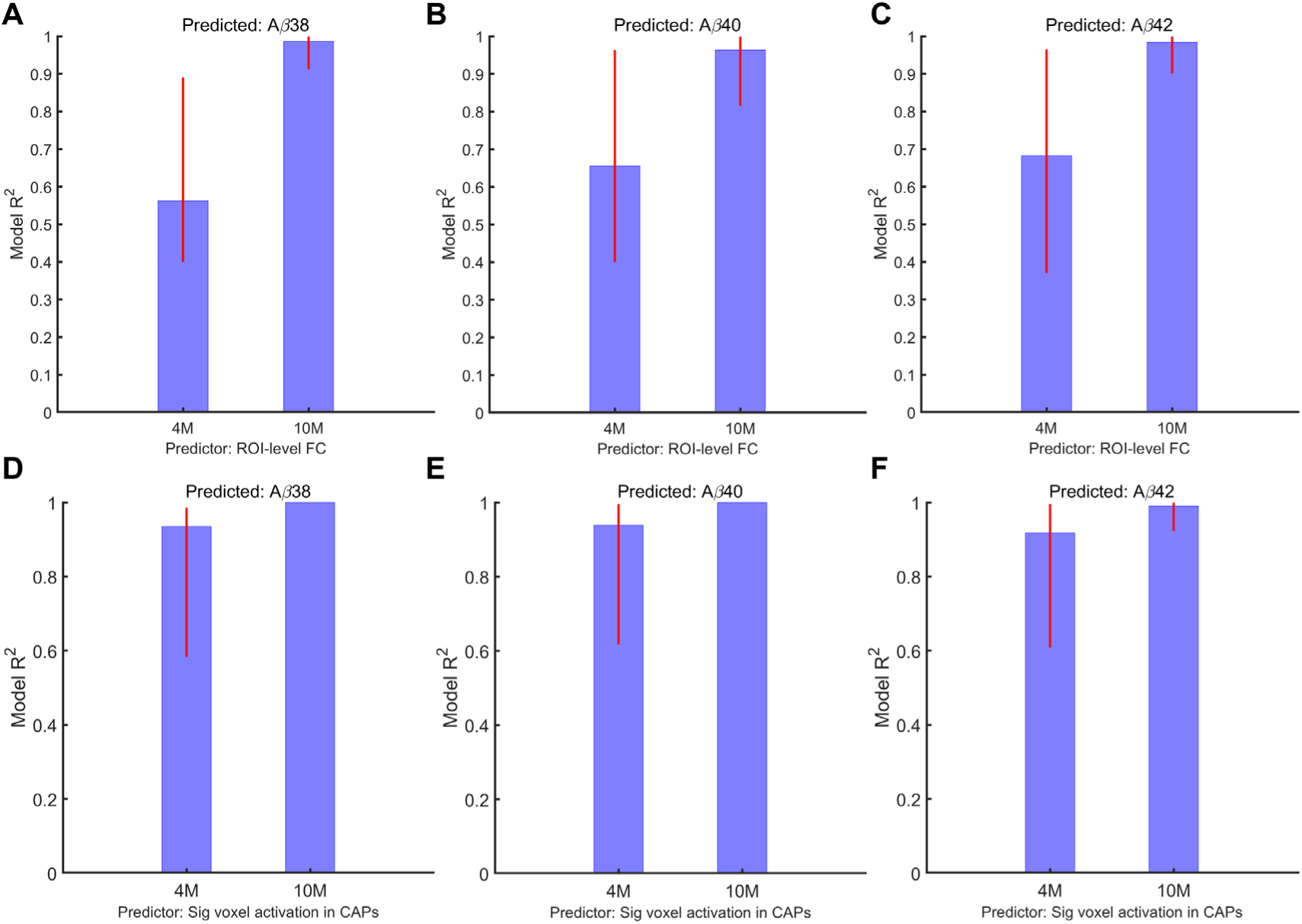
PLSR model *R*^2^ for predicting the BBMs at the plaque stage using ROI-based FC (top panels) and CAPs (bottom panels) at the pre-plaque (4M) and plaque (10M) stages. Here we used pairwise, WT-normalized FC values between all 34 ROIs considered in Figure 3 A-B (**A, B, C**), and voxel-wise, WT-normalized, one-sample T-test statistic values in each TG subject (**D, E, F**) for the voxels found to be significantly (de)activated in the combined CAPs one-sample T-test statistic maps (Figure 4 A, D, G, J, M, P) to explain the WT-normalized values of *Aβ*38, *Aβ*40, and *Aβ*42 in each individual TG subject respectively. Red vertical lines: FWER-corrected 95% confidence intervals.

We next sought the functional connections (for ROI-FC) and voxel-clusters (for CAPs) that contributed most prominently to the high model *R*^2^ at both 4M and 10M. Specifically, we calculated the regression weights for all measures from each fitted model and then identified the measures for which the FWER corrected 95% confidence intervals for the weights did not cross zero, implying a significant positive or negative effect on the amyloid levels in blood. We flipped the signs of weights for functional measures for which the TG animals had lower mean value than WT animals. This ensured the interpretation that positive and negative weights imply higher amyloid levels in TG animals correspond to greater and lower severity, respectively, in the explanatory measure. Figure 6A shows the ROI-pairs with highest 20%, significant positive weights at 4M and 10M, whose FC values explained the variance in the levels of *Aβ*42. At 4M, CA3-Lim, and several connections between the DMLN and the LCN networks while at 10M, FC between CA3 and CA1, Cg, Aud, MS as well as BFB-Ent were the main contributors to the explanatory power of the model. Also, the magnitude of weights, in general, was higher at 10M than at 4M.

**Figure 6:**
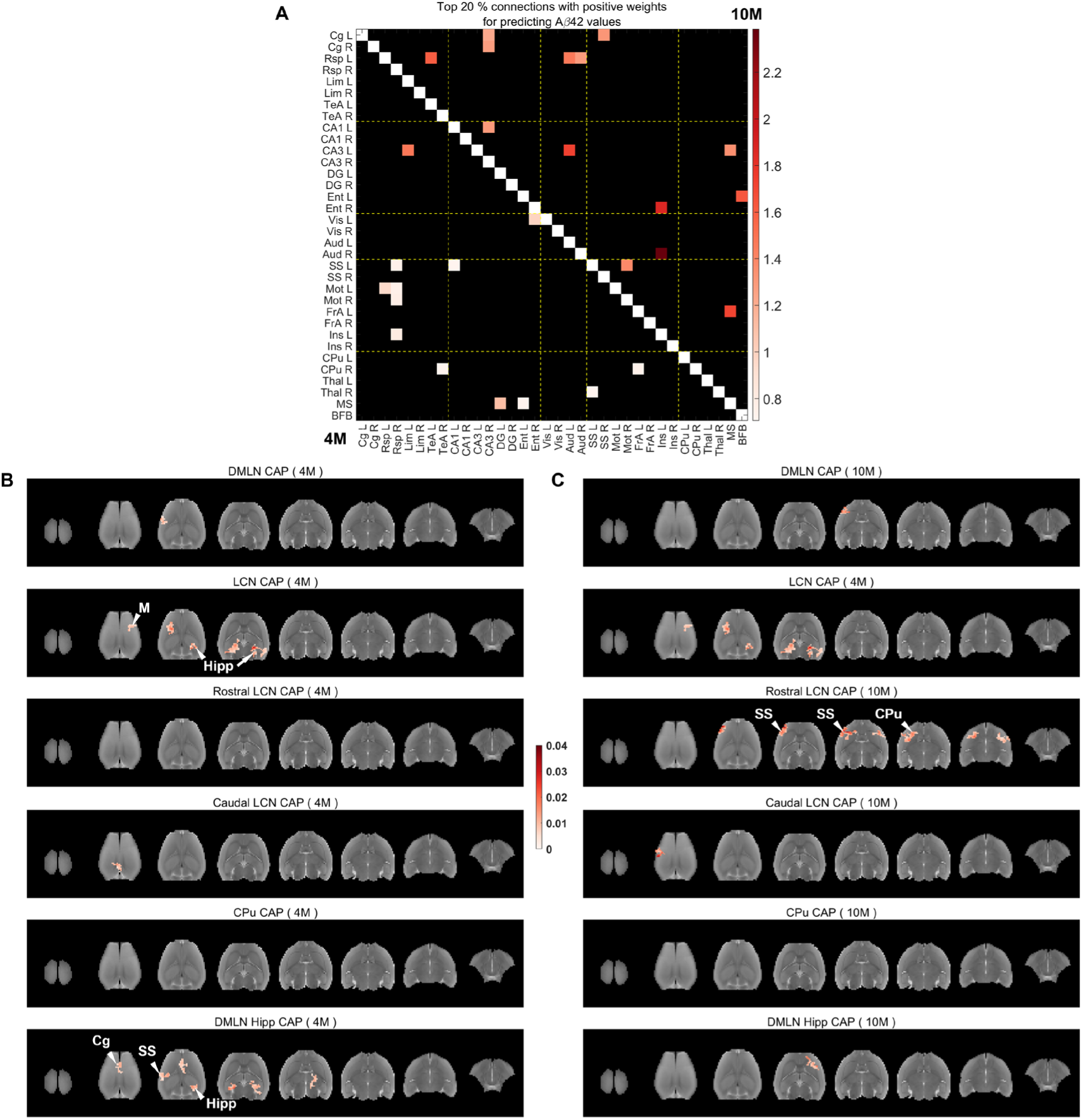
Salient functional connections (A) and voxel-clusters (B-C) within CAPs at pre-plaque and plaque stages that explain the *Aβ*42 values in the TG animals. **A:** Top 20% highest positive beta weights of functional connections with significant 99% confidence intervals at 4M (below the diagonal) and 10M (above the diagonal). **B, C:** Positive beta weights for salient clustered voxels (cluster-size: 20 voxels, in plane), for which FWER corrected 95% confidence intervals do not cross zero, in each CAP at 4M (B) and 10M (C).

Figures 6B and 6C show the salient voxel-clusters from each of the six CAPs with significant positive weights for explaining the levels of *Aβ*42 at 4M and 10M respectively. At 4M, prominent voxel clusters were found in the DMLN-Hipp CAP in the Hipp, Cg, and SS and, in the LCN CAP, in the Hipp and Mot regions. At 10M, CPu, SS in the DMLN-Hipp CAP, and CPu, SS, Ins in the Rostral LCN CAP showed prominent voxel clusters.

Supplementary Figures 4 and 5 show the prominent FC and voxel-clusters in CAPs with significant positive weights for explaining levels of *Aβ*38, and *Aβ*40 respectively. The explanator ROI-pairs for *Aβ*40 values were very similar to those for *Aβ*42 at 4M; at 10M, in addition to CA3-Aud and CA3-MS which also explain *Aβ*42 levels, MS-SS, and MS-FrA were specific for *Aβ*40 (Supplementary Figure 5A). In the DMLN-Hipp CAP, CPu and ventral Hipp at 4M and 10M, respectively, display voxel clusters explaining *Aβ*40 levels, in addition to those found relevant for *Aβ*42 (Supplementary Figure 5B-C). In the LCN CAP, Mot, SS, Hipp, Th, and CPu show clusters whose activations at 4M explain the *Aβ*40 levels in TG animals (Supplementary Figure 5B). Cg-Aud FC at 4M while FC of Ent with Ins and Cg, and of Mot with CA1 and DG at 10M uniquely explain the *Aβ*38 levels (Supplementary Figure 4A). Voxel clusters in SS in all 4M CAPs except the Rostral LCN CAP, while, at 10M, very localised clusters in the CPu in the Rostral LCN CAP, BFB in the Caudal LCN CAP, and ventral Hipp in the DMLN-Hipp CAP prominently explain the *Aβ*38 levels in the TG animals. Supplementary Figures 6-8 show the top 20% ROI-pairs and prominent voxel-clusters that show significant negative regression weights in the PLSR models that explain the levels of *Aβ*38, *Aβ*40, and *Aβ*42 in the TG animals, respectively.

### Relationship between functional changes, blood-based markers and memory alterations

Finally, we assessed whether the BBMs and/or RS-functional measures changes could explain the alterations we observed in the memory performance tasks in the TG animals. We focused on reference and working memory errors, two of the nine behavioral performance measures acquired on day 10 of the training (Figure 2), as they showed a significant genotypic groups difference. Their WT-normalized values were used as response variables for fitting PLSR models using WT-normalized BBMs, ROI-based FC or activations of significant voxels in CAPs. Amyloid markers showed poor prediction of both behavioral measures (model 𝑅𝑅^2^∼ 0.2, Supplementary Figure 9). ROI-based, WT-normalized FC values at 4M and 10M showed a model variance 𝑅𝑅^2^∼ 0.6 in explaining reference memory errors (Figure 7A). For working memory errors, 4M FC values had a model 𝑅𝑅^2^∼ 0.6 while the 10M FC values showed high explanatory power but with very long 95% confidence intervals that included 0 indicating not a robust fit for the model (Figure 7B). Spatial properties of CAPs at both 4M and 10M, however, showed a high model 𝑅𝑅^2^(∼ 0.8 − 1) for both behavioral measures (Figure 7 C-D). Interestingly, spatial properties of CAPs at 4M explained more variance in both behavioral markers than at 10M, suggesting the prognostic value of CAPs.

**Figure 7:**
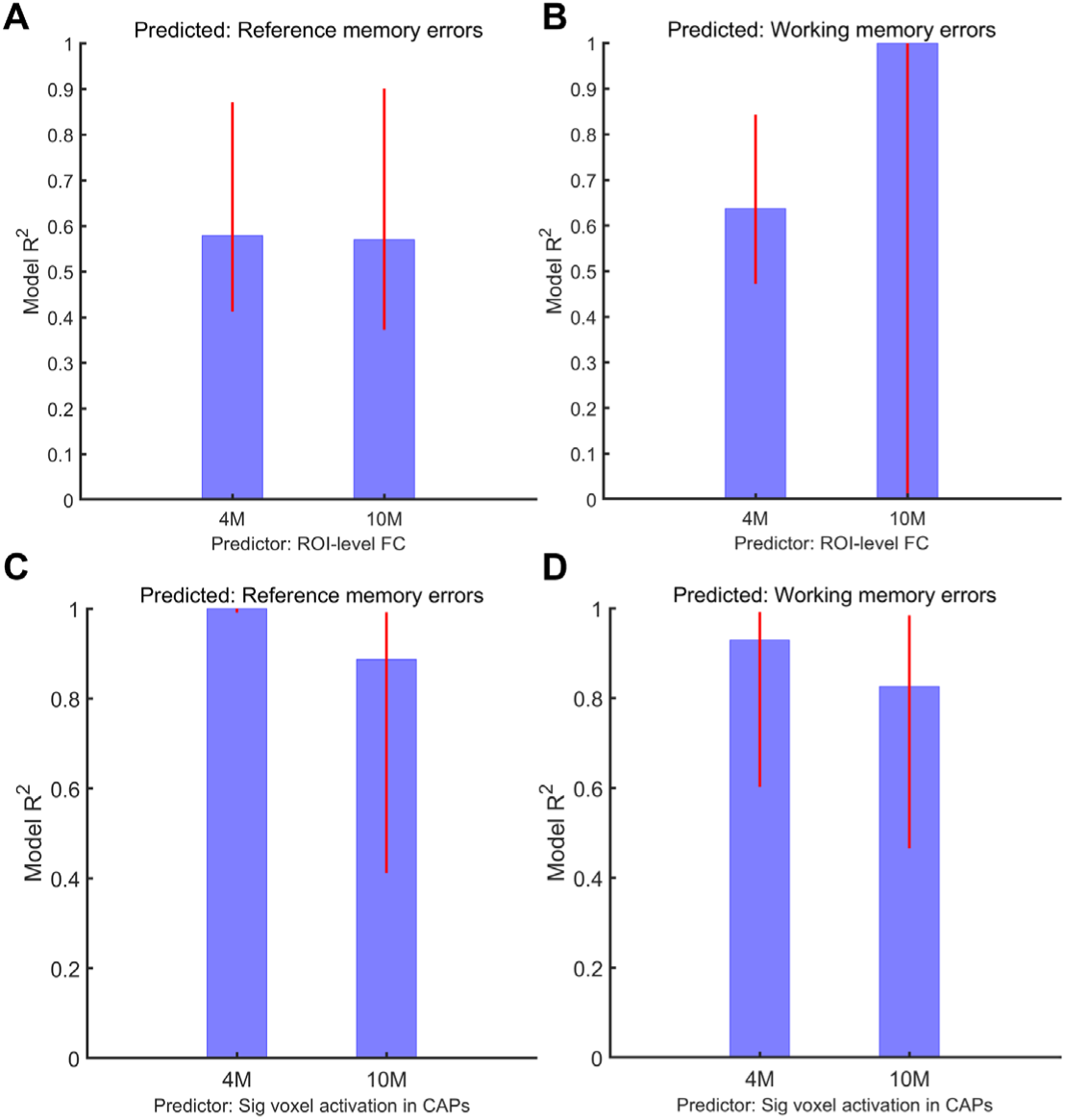
PLSR model *R*^2^ for predicting memory tasks performance (reference and working memory errors) on acquisition day 10, measured during the plaque stage, using ROI-based FC (top panels) and CAP spatial properties (bottom panels) at the pre-plaque (4M) and plaque (10M) stages. Here, we used pairwise WT-normalized FC values between all 34 ROIs considered in Figure 3A-B (**A, B**) and voxel-wise, WT-normalized one-sample T-test statistic values for each TG subject (**C, D**) for the voxels found to be significantly (de)activated in the combined CAP one-sample T-test statistic maps (Figure 4 A, D, G, J, M, P) to explain the WT-normalized values of reference memory errors and working memory errors in each individual TG subject respectively. Red vertical lines: FWER -corrected 95% confidence intervals.

Next, we identified the salient functional measures at 4M and 10M that most prominently explain the memory impairments. Figures 8 and 9 show the significant and positive weights for the most relevant functional measures (FC and CAPs) for explaining the reference and working memory impairments, respectively, while Supplementary Figures 10 and 11 display the significant and most prominent negative weights for the fitted PLSR models. Reference memory errors in the TG animals are explained most prominently by FC changes for SS-Ins and FrA-SS connections at 4M, and BFB-Mot, CA1-Ent, and inter-hemispheric Rsp connection at 10M (Figure 9A). FrA in the LCN and Rostral LCN CAPs and SS in the DMLN-Hipp CAP at 4M while, at 10M, Rsp in the Rostral LCN, the Caudal LCN, and the DMLN Hipp CAPs, and Vis in the DMLN and the LCN CAPs show most explanatory voxel clusters with positive weights for reference memory errors (Figure 9 B-C). In the case of working memory errors, FC between Ent and Cg, Vis and between SS and TeA at 4M while FC of Aud with CA1 and Ins, of Ent with Vis and Thal, and between Thal and Ins, explain the most variance across TG animals (Figure 10A). Most explanatory voxels clusters for working memory errors were found in the Rsp in the Caudal LCN and the CPu CAPs, in Vis in the DMLN CAP, and in SS in the DMLN Hipp CAP at 4M while, at 10M, in the BFB in the Rostral LCN and CPu CAPs (Figure 10B-C).

**Figure 8:**
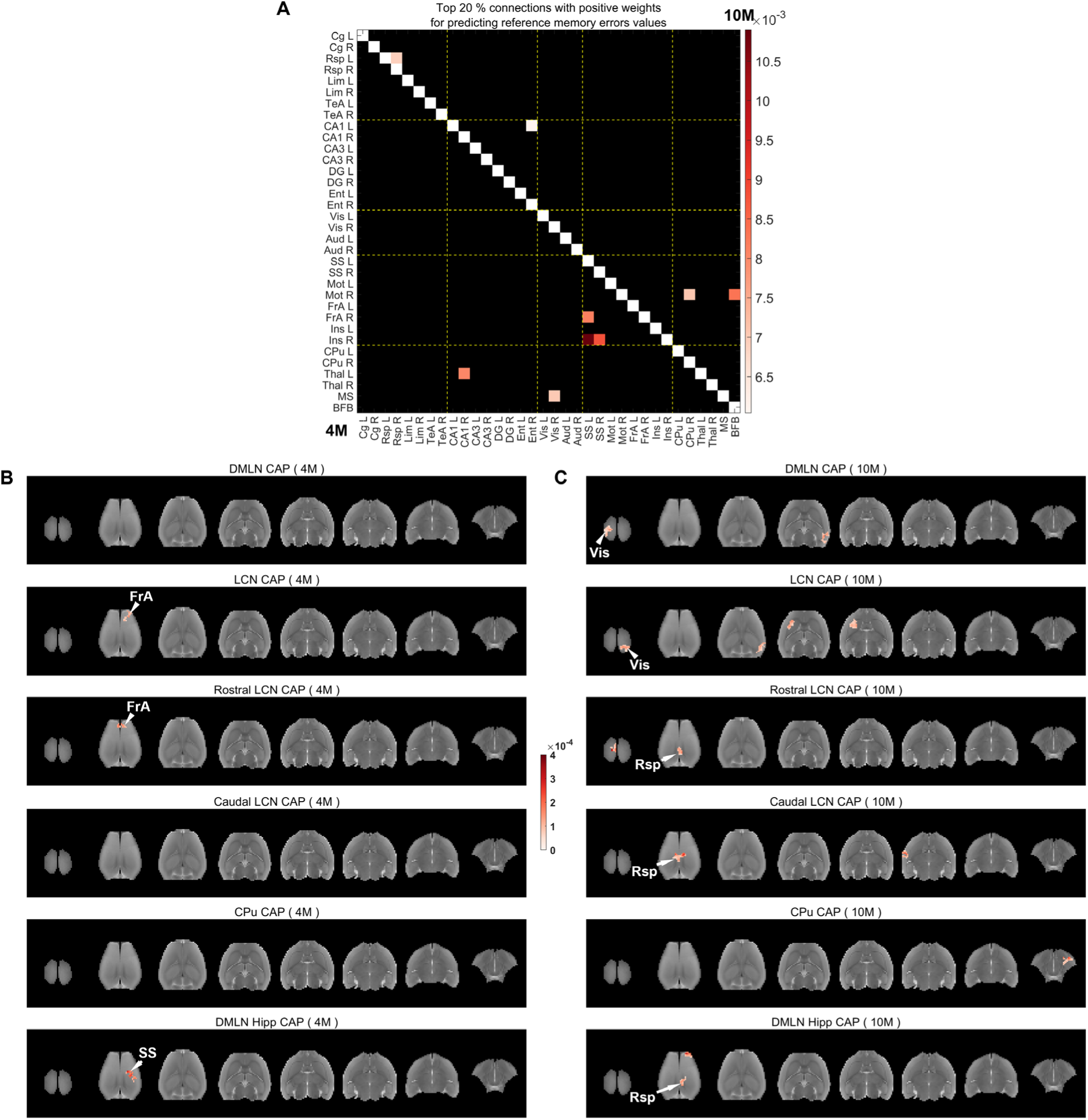
Salient functional connections (A) and voxel-clusters (B-C) within CAPs with positive beta weights at pre-plaque and plaque stages that explain WT-normalized reference memory errors in the TG animals. **A:** Top 20% highest positive beta weights of functional connections with significant 99% confidence intervals at 4M (below the diagonal) and 10M (above the diagonal). **B, C:** Positive beta weights for salient clustered voxels (cluster-size: 20 voxels, in plane), for which FWER corrected 95% confidence intervals do not cross zero, in each CAP at 4M (B) and 10M (C).

**Figure 9:**
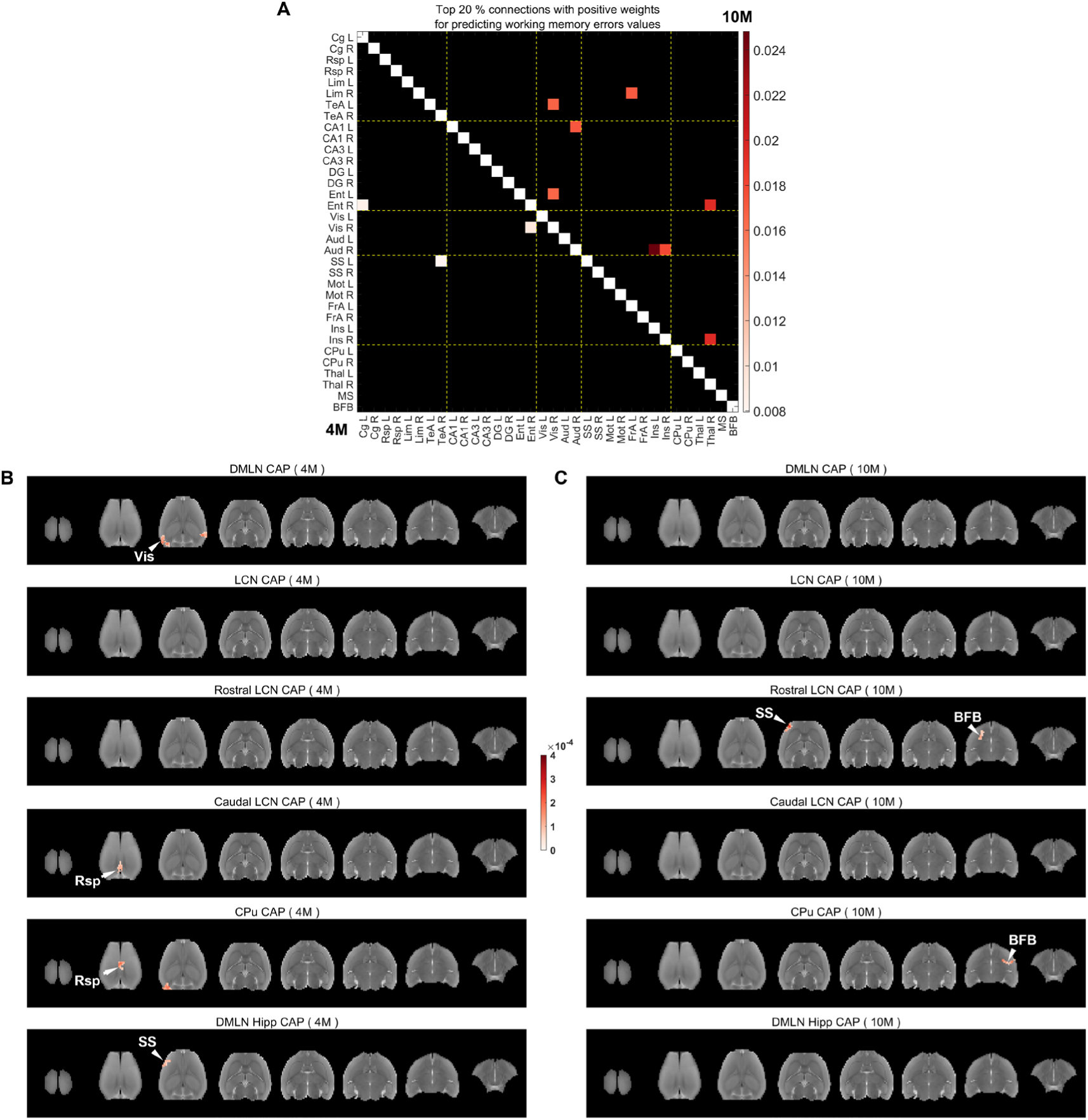
Salient functional connections (A) and voxel-clusters (B-C) within CAPs with positive beta weights at pre-plaque and plaque stages that explain WT-normalized memory errors in the TG animals. **A:** Top 20% highest positive beta weights of functional connections with significant 99% confidence intervals at 4M (below the diagonal) and 10M (above the diagonal). **B, C:** Positive beta weights for salient clustered voxels (cluster-size: 20 voxels, in plane), for which FWER corrected 95% confidence intervals do not cross zero, in each CAP at 4M (B) and 10M (C).

## Discussion

In this study, we aimed to characterize, in the TgF344-AD model, phenotypic markers of Alzheimer’s disease in blood plasma, resting-state functional connectivity and dynamics, as well as memory impairments and investigate their inter-relationships. At the plaque stage, we found significantly elevated levels for amyloid markers (*Aβ*38, *Aβ*40, *Aβ*42), working and reference memory impairments, a reduced magnitude of RS-FC between the hippocampus and the default-mode-like network as well as the lateral cortical network and an overall hypoactivation in several regions in all six co-activation patterns in the TG animals vis-à-vis the WT. At the pre-plaque stage, while the RS-FC did not exhibit any alterations, several CAPs showed hyperactivation, especially in the DMLN and hippocampal regions, in the TG group. Compared to FC, CAPs’ spatial properties explained higher variance in both the BBMs, and memory impairments. Most interestingly, properties of pre-plaque CAPs explained more variance in the memory impairments than at plaque stage suggesting prognostic relevance of brain’s transient functional states.

The levels Aβ40 and Aβ42 found in the blood plasma of TgF344-AD animals at 11 months of age were similar to those reported at 8-12 months of age in this model^16^. Immunohistochemistry analysis in our previous study in the TgF344-AD model demonstrated the presence of amyloid plaques in the hippocampus and retrosplenial cortex at 6 months^17^. Therefore, the elevated plasma levels of all three amyloid markers in the TG animals imply that there is production of Aβ, and given the histological evidence from previous studies, indicate widespread presence of amyloid plaques in the brain of these model rats. We also tried to obtain the plasma levels of GFAP; however, they were below the detection threshold in all animals. On the other hand, there was no significant difference in the levels of neurofilament light chain (NfL) levels between the TG animals and the WT implying no significant axonal degeneration in this model at this age.

Memory assessments in the TgF344-AD model have predominantly used the Morris water maze (MWM) that measures spatial memory alterations. In our study, we mainly looked at reference and working memory alterations using the radial arm maze. Our findings demonstrate very clearly that over the acquisition period WT rats progressively performed the task better compared to TG rats. Parameters such as total number of entries, total travelled distance, and most importantly working memory errors showed a clear learning-effect over time in the WT animals but not in the TG resulting in the genotypic group difference towards the end of the acquisition period. Other parameters such as total duration, time spent and distance travelled in non-baited arms, and reference memory errors showed an overall impairment in the TG animals vis-à-vis the WT, but not a progressive deterioration. A recent study that assessed reference and working memory using water radial arm maze in female TgF344-AD rats found significantly higher reference and working memory errors at 12 months of age but no difference with the WT at 6 or 9 months of age^18^.

Our assessment of functional network connectivity using RS-fMRI revealed hippocampus’ correlation and anticorrelation with the DMLN and the LCN, respectively, was reduced in the TG animals but only at the plaque stage. Reduced functional correlation between the hippocampus and DMN regions has been reported in AD patients as well as in MCI patients^10,19–21^.. The LCN is the rodent analog^22^ of human task positive network (TPN), that is usually anticorrelated to the DMN^23^. The DMN-LCN anticorrelation has been found to be reduced in AD^24^. We did not find any significant difference between the network-level connectivity at the pre-plaque stage. Moreover, the ROI-level FC was not found to be altered between the genotypes at either of the ages, although we found an age-dependent increase of correlation and anticorrelation of several Hipp ROIs with the DMLN and the LCN ROIs, respectively, in the WT group alone.

Temporal fluctuations in the FC, ignored by static FC, could be informative and sensitive to measure alterations in the TG animals^13,14^. We captured these temporal fluctuations using transient functional brain states in the form of co-activation patterns and found a clear dichotomous picture of changes in spatial co-activations. At the pre-plaque stage, most CAPs showed hyper(de)activation in especially the DMLN, LCN and hippocampus regions while a large-scale hypo(de)activation was observed at the plaque stage. The concept of hypersynchronous network activity playing a significant role in the development of AD has gained increasing acceptance in recent years^25^. It is now widely accepted that abnormally high neuronal network activities precede the formation of amyloid plaques in animal and cell models^25–27^. In children at risk for autosomal dominant AD carrying the presenilin 1 mutation, FC of the posterior cingulate cortex with medial temporal lobe regions was found to be increased compared to non-carrying children^28^. During memory encoding tasks, hippocampal hyperactivation and decreased inactivation of the hub regions within the DMN have been reported to be a consistent fMRI signature of AD, particularly at the early stages^25,26,29^. The role of this hyperactivation could be compensatory^29^ but growing evidence that individuals with AD exhibit increased susceptibility to seizures^25^ suggests a pathogenic one. Cognitive decline in individuals with AD and epilepsy tends to occur earlier, and antiepileptic drugs that can reduce neuronal hyperexcitability have been found to enhance memory performance in MCI patients^25^. The hyper(de)activations found in CAPs is also in line with our finding in a mouse model of HD at 3 months of age^14^. The hypo(de)activation found in CAPs at the plaque stage can explain the reduced magnitudes of hippocampal – DMLN/LCN FC observed in these animals. This observation is also consistent with robust findings in patients of AD at symptomatic stages^10^.

Characterizing the pathological, behavioral, and resting-state functional phenotypes in the same TgF344-AD model rats allowed us to investigate inter-phenotypic relationships. In our previous publications, we have performed cross-validated classification using RS functional measures like FC and CAPs and shown that spatial properties of CAPs, in particular, can accurately distinguish animals of transgenic models of neurodegenerative disorders from the WT^13–15^. Here we inquired whether changes in pre- and plaque stage FC and CAPs in an individual TG animal can accurately explain the alterations in plasma amyloid and memory performance levels at the plaque stage in the same animal. We found that blood plasma amyloid levels were well explained by both the ROI-based FC and CAPs at both stages. However, higher *R*^2^ for the plaque stage FC and CAP changes imply amyloid levels in blood plasma when amyloid plaques are present in the brain are explained better by the functional changes at the same stage than the functional changes when only soluble amyloid, but no plaques are present in the brain of TG rats. Preferential accumulation of Aβ in the DMN has been linked to the reduced DMN connectivity both in non-demented old individuals^30^ as well as in AD patients^31,32^. We found that the three plasma amyloid marker values, especially Aβ40 and Aβ42, were explained by similar prominent functional connections and voxel-clusters in CAPs at the plaque stage. FC between hippocampal, DMLN regions as well as the medial septum and basal forebrain while voxel-clusters in the basal forebrain of the Rostral LCN and CPu CAPs at the plaque stage were salient in explaining the amyloid markers in the blood plasma. Interestingly, unlike FC, pre-plaque stage CAPs’ activations were also very accurate (*R*^2^∼0.9) in explaining the plasma amyloid markers and prominent contributors were found in the DMLN and hippocampal regions in the DMLN-Hipp CAP.

When it comes to the memory impairments, we found that the RS-CAPs explained them with higher *R*^2^ values than RS-FC in general. RS-FC has been used to predict cognitive impairment in AD assessment scale in patients using partial least squares regression^33^. A recent study also used connectome-based predictive modeling to predict both the tau deposition as well as cognitive scores for memory and mental state using RS-FC in an autosomal dominant AD cohort of Presenilin-1 E280A carriers’ mutation^34^. Most recently, Fadel et al. also used machine learning approach to longitudinally investigate the relationship between RS-FC, obtained in awake APP/PS1 model mice, and learning and memory^35^. However, no study has investigated the relationship between cognitive impairments and measures of dynamic FC in AD. In this context, our finding that RS-CAPs are more informative than FC for explaining memory impairments in this translationally relevant model stands out. Most interestingly, CAPs at pre-plaque stage were more informative than those at the plaque stage for explaining the reference and working memory errors in the TG animals at plaque stage. Reference memory errors were explained by localized voxel-clusters in the frontal association cortex in pre-plaque stage CAPs and in the retrosplenial cortex in plaque stage CAPs. On the other hand, localized voxels in the retrosplenial cortex from pre-plaque stage CAPs were important contributors for explaining the working memory errors. Frontal association cortex is important for episodic memory for novel object recognition in rats^36^. Retrosplenial cortex, a key component of the DMLN, is densely connected with the hippocampus and is involved in memory processing in rats^37,38^. As resting-state networks have a high degree of similarity to functional networks found in tasks^39^, our findings demonstrate that changes in coactivation of regions, typically involved in memory processing, in dynamic functional configurations of the resting-state are salient markers of memory impairments in this translational model of AD. Finally, the importance of pre-plaque stage CAPs for explaining memory impairments underscores their prognostic potential.

### Limitations of the study

One of the limitations of our study is that only male rats were used to limit the variability with the sample size determined based on power calculations. A more severe amyloid pathology and neuroinflammation but less severe cognitive deficits have been reported in female TgF344-AD rats as compared to males^40,41^. Therefore, rats of both sexes are included in all our future studies. Radial arm maze used to assess reference and working memory relies on task motivation which was achieved by food restriction that can cause stress and anxiety in animals and act as a confound. Also, rats can use olfactory clues and egocentric navigation strategies instead of purely allocentric ones.

The influence of olfactory clues was minimized by cleaning the RAM with a product that had a strong, penetrating odor. To avoid egocentric navigation, rats were placed in the maze in a random orientation at the start of the trial. CAPs were obtained by clustering the concatenated, preprocessed images from animals of both genotypic groups at both ages as we were interested in investigating both the genotype and age effects. This approach limits identification of age and genotype specific CAPs. Finally, we did not use partial least square regression to do an out-of-sample prediction, as we were mainly interested in explaining the amyloid markers or memory impairments using RS-fMRI measures. It would be straightforward to make cross-validated predictions of outcomes of left-out subjects to test the predictive capacity of RS-FC and CAPs.

## Materials and Methods

### Ethical statement

All procedures were performed in strict accordance with the European Directive 2010/63/EU on the protection of animals used for scientific purposes. The protocols were approved by the Committee on Animal Care and Use at the University of Antwerp, Belgium (permit number 2021-52) and all efforts were made to minimize animal suffering. The reporting in the manuscript follows the recommendations in ARRIVE guidelines.

### Animals

Fifteen male TgF344-AD rats and 15 male wild type Fischer rats were used. Animals were bred in-house, initiated with breeding pairs that were bought from the rat resource and research center (RRRC). Rats were subjected to MRI acquisition at the age of 4 and 10 months. At 4 months of age, only soluble amyloid but no plaques are observed while at 10 months amyloid plaques are ubiquitous in the brain of TgF344-AD rats^17,42^. Spatial memory deficits are observed from 6 months of age^43^. Behavioral evaluation was performed in 10.5-11 months old rats after the final MRI scanning session. All rats were group-housed with two animals per cage. Room temperature and relative humidity were controlled, and *ad libitum* food and water were present, except during the behavior testing phase (see behavior data acquisition section). Two animals (1 WT and 1 TG rat) reached predefined humane endpoints before the age of 10 months and were euthanized. As a result, MRI data could only be acquired for these animals at 4 months of age.

### Behavior data acquisition

A custom-built eight-arm radial arm maze was used to evaluate visuospatial learning and memory in the 10.5-11-month-old rats. The maze comprised a central octagonal area with sides of 9.8 cm and eight evenly spaced arms extending outwards, each 48 cm long, 8.3 cm wide, and enclosed by 25.5 cm high walls. Bird spikes were installed along the maze edges to prevent escape while maintaining optimal visibility of the surroundings. The setup was surrounded by both two-dimensional and three-dimensional visual cues. During trials, the researcher was concealed out of view from the rats. Tests were performed in a dimly red-lit room supplemented by an infrared light (Stoelting Europe, Ireland). Tracking was conducted live via ANY-maze software (Stoelting Europe, Ireland) using an overhead infrared USB camera (Stoelting Europe, Ireland).

The rewards consisted of halved honey pops. To provide equal olfactory stimulation across all arms, inaccessible rewards underneath a mesh cover were always present in cups at the end of each arm. Depending on the testing phase, some or all arms contained accessible rewards placed below floor level to avoid visibility from the center of the maze. The maze was cleaned with 70% propanol (IPASEPT 70, VWR Chemicals, Belgium) between trials to eliminate odor traces.

The testing protocol consisted of three phases: familiarization, habituation and acquisition training. The familiarization phase lasted seven consecutive days, during which rats were familiarized with the rewards in their home cages and with handling by the training personnel. During the habituation phase, spanning five to seven days, rats were exposed to the maze daily and the rewards were made accessible in all arms. Initially, rewards were visibly placed at the end of each arm and then hidden below floor level from the third session onward. Each day consisted of a single trial, lasting a maximum of 15 minutes or until all rewards had been found and eaten. If rats located at least five out of eight rewards by day five, they proceeded to the next phase; otherwise, habituation continued for up to two additional days. The acquisition phase lasted ten consecutive days, during which rewards were accessible in a fixed set of four arms. Each acquisition training day consisted of three trials of up to five minutes each, with an interval of 30-50 min.

For every trial, rats were placed in the center of the maze with a random orientation to promote allocentric over egocentric learning. From the start of the habituation phase, food restriction was employed to motivate exploration. Rats received 10 g of standard food pellets on testing days (in addition to rewards found in the maze) and 12 g on non-testing days, corresponding to a 40% caloric restriction.

### Analysis of behavior data

During acquisition period, the animal behavior was tracked by ANY-maze software that measured several parameters: total trial time, total distance travelled, number of entries into each arm, the number of arms that were entered, the total number of entries, the number of mistakes, and average speed. Additionally, we calculated metrics such as percentage of reference memory errors (percentage of entries into non-baited arms), percentage of distance and time spent in non-baited arms, latency to first entry into baited and non-baited arms), and percentage of working memory errors (percentage of re-entries into already entered arms). For each acquisition training day, average values across the three trials were used for further statistical analysis.

### Serum samples acquisition and analysis

Serum samples were acquired by lateral tail vein catheterization from all animals at 11 months of age at the time of the MRI scans. They were analyzed for Neurofilament Light (NfL), Glial Fibrillary Acidic Protein (GFAP) and different Amyloid bèta forms, namely Aβ38, Aβ40 and Aβ42, using the R-PLEX Human Neurofilament L (K1517X-2), R-PLEX Human GFAP (K1511MR-2) and Aβ Peptide Panel 1 6E10 (K15200E-1) assays respectively, following the manufacturer’s protocol. Briefly, samples were incubated undiluted on the pre-coated plate. After incubation at room temperature, the wells were washed and incubated with SULFO-TAG labeled detection antibody. Afterwards, the plate was washed, reading buffer was added and reading was performed using the MESO QuickPlex SQ 120MM. Concentration calculations were calculated using Discovery Workbench v4.0 software from MesoScale Discovery. Further statistical analysis was performed by Graphpad Prism

### MRI data acquisition

This manuscript uses the 3D anatomical and RS-fMRI data which are part of a larger study in which, following the 3D anatomical scan, a series of fMRI acquisitions were performed in the following order: BOLD-RS-fMRI, visual stimulus-based BOLD-fMRI, CBV-weighted RS-fMRI, and visual stimulus-based CBV-weighted fMRI.

All data were acquired on a 9.4 T Biospec MRI system (Bruker BioSpin, Germany) equipped with a 2×2 array rat head receiver RF coil, using Paravision 360 software. The rats’ heads were immobilized using an MR-compatible stereotaxic device with blunt earplugs and a tooth bar. A cannula was placed in the tail vein for intravenous delivery of ultrasmall superparamagnetic iron oxide (USPIO) nanoparticles, a T2 relaxation-lowering MRI contrast agent. A fibre-optic coupled to a white LED was placed centrally in front of the animal for binocular visual stimulation.

### Anesthesia

Prior to the scan, anesthesia was induced using isoflurane (5%, Forene; Abbott, Belgium) in a 30% O₂ and 70% N₂ mixture, followed by subcutaneous (s.c.) administration of a 0.05 mg/kg bolus of medetomidine hydrochloride (Domitor, Pfizer, Germany). After bolus administration, isoflurane concentration was progressively reduced to 0.4% during fMRI acquisition. Fifteen minutes post-bolus injection, a continuous s.c. infusion of 0.1 mg/kg/h medetomidine was initiated and sustained for the entire scan duration. At the end of the session, the reversal agent atipamezole (0.25 mg/kg, Antisedan, Pfizer, Germany) was administered s.c.. Physiological parameters, including respiration rate and blood oxygenation, were monitored using a pressure-sensitive pad and a pulse oximeter (Small Animal Monitoring and Gating System, SA Instruments, Inc., USA). Body temperature was maintained at (37.0 ± 0.5)°C using a feedback-controlled warm air system (MR-compatible Small Animal Heating System, SA Instruments, Inc., USA).

### fMRI parameters

Slice positioning for RS-fMRI was standardized using orthogonal multi-slice T2-weighted TurboRARE images to ensure consistency across subjects. A B_0_-field map was acquired during each session, followed by localized shimming to optimize field homogeneity within an ellipsoid volume covering the brain.

A 3D T2-weighted TurboRARE anatomical scan covered the entire brain with a spatial resolution of (0.137 × 0.137 × 0.3) mm³ and the following parameters: repetition time (TR) 1800 ms, echo time (TE) 11 ms (TE_eff_ 44 ms), RARE factor 16. The field of view (FOV) was (35 × 35 × 16) mm³ with an acquisition matrix of [256 × 256 × 53], and the scan duration was 14 min 24 s.

RS-fMRI was performed at 35 min post-medetomidine bolus injection using a T2*-weighted EPI sequence (TR = 600 ms, TE = 18 ms, 20 slices, 1 mm slice thickness, 0.1 mm gap). Images were acquired with a (35 × 35) mm² FOV and a matrix size of [96 × 96], yielding voxel dimensions of (0.364 × 0.364 × 1) mm³. A total of 2000 frames were acquired continuously: the first 1000 were collected during resting state (10 min; 35-45 min post-medetomidine bolus), followed by an additional 1000 frames during a visual stimulation paradigm (10 min; 45-55 min post-medetomidine bolus)..

Approximately 55 minutes after medetomidine bolus administration, 10 mg/kg USPIO nanoparticles (Molday ION, BioPal, Worcester, USA) were injected via the tail vein catheter using an MRI-compatible infusion pump. CBV-weighted fMRI data were acquired using the same acquisition protocol as for the BOLD fMRI, with the exception of a shorter TE of 10 ms. 2000 volumes were acquired of which 1000 in resting state and 1000 upon visual stimulation.

For one WT animal, we observed a darker signal distribution already in the BOLD RS-fMRI, indicative for leakage of the USPIO nanoparticles via the tail vein catheter before the BOLD-RS-fMRI acquisition. As this would give a mixed opposite BOLD and CBV weighted effect on the signal, the RS-fMRI data for this animal were excluded from both ages from further analysis.

### Pre-processing of BOLD-RS-fMRI data

Pre-processing of the RS-fMRI data, including realignment, normalization and smoothing, was performed using SPM12 software (Statistical Parametric Mapping, http://www.fil.ion.ucl.ac.uk). The RS-fMRI EPI images were realigned to the first EPI image using a 6-parameter rigid body spatial transformation estimated using a least-squares approach. After realigning the RS-fMRI images, the translational and rotational realignment parameters were examined. For one WT animal at the 4-month timepoint, the translational values exhibited excessive movement, indicating poor image quality. As a result, this animal was excluded from further analysis. Then, both the EPI-images and the subject-specific 3D images were debiased and were co-registered to the debiased brain-masked subject-specific 3D image using the old-normalise function in SPM12, which comprises the estimation of an affine transformation followed by the estimation of a non-linear deformations. Afterwards, the old-normalise function in SPM12 was used to normalize the debiased subject-specific 3D images to a study-specific 3D template, created from the 3D MRI data of all subjects at both ages using Advanced Normalization Tools (ANTs). Finally, realigned EPI images were normalized to the study-specific 3D template by applying the combined transformations. The normalized EPI images were smoothed in-plane using a Gaussian kernel with full width at half maximum (FWHM) of twice the pixel size (FWHM [0.7291×0.7291×1.1] mm^3^).

Images were then filtered voxel-wise using a 0.01-0.2 Hz Butterworth band-pass filter. Finally, quadratic detrending was performed on the filtered images using in-house MATLAB scripts. Ten frames at the start and the end of the scan were excluded before and after filtering to avoid transient effects and to minimize boundary effects of filtering respectively.

### Functional connectivity analysis

For the ROI-based and network-based FC analysis of each subject, Global signal regression was performed on preprocessed RS-fMRI scans. Next, we extracted the ROI-based BOLD signals of 34 ROIs (Figure 3) in total belonging to 5 resting-state networks:

1. The default-mode-like network (DMLN): cingulate (Cg), retrosplenial (Rsp), limbic (Lim), and temporal association (TeA) cortices.
2. The hippocampal network (Hipp): CA1, CA3, dentate gyrus (DG), and entorhinal (Ent) cortex.
3. The sensory network (Sens): visual (Vis) and auditory (Aud) cortices.
4. The lateral cortical network (LCN): the somatosensory (SS), motor (M), frontal association (FrA), insular (Ins) cortices.
5. The subcortical network: caudate putamen (CPu), thalamus (Thal), medial septum (MS), and basal forebrain (BFB).

We first estimated the transformation of the 3D template to the Fischer 344 rat brain atlas, then used the inverse transformation to warp the atlas-based label fields of these ROIs to the template space and obtained BOLD time-course for each ROI by averaging BOLD signals across all voxels belonging to it.

ROI-based FC matrix for each subject were obtained by calculating the Pearson’s correlation between BOLD time courses of each pair of ROIs. Subsequently, we obtained the network-based FC matrix by averaging FC values across all pairs of ROIs belonging either to the same network (diagonal values, within network FC) or between different networks (off-diagonal values, between network FC).

### Co-activation Patterns (CAP) analysis

For the CAP analysis, we regressed out the ventricular and white-matter signal from the filtered and detrended BOLD signal of each voxel. The resulting signal for each voxel was then z-scored so that it had zero mean and unit standard deviation. We then applied a whole-brain mask– excluding the olfactory bulb and the cerebellum - to the preprocessed images of each subject and concatenated these from all subjects from both genotypic groups and ages to create a combined image series. The image series was then transformed into an n x m -dimensional matrix, where n was the total number of frames from all subjects, and m was the total number of voxels in the brain mask. We retained the signal values of the voxels that had the top 10% highest and bottom 5% lowest z-scored signal intensities in each frame while setting the rest to zero and clustered all time frames using a K-means++ algorithm by assessing the correlation distance between every pair of m-dimensional vectors. The algorithm partitions the n vectors (time frames) into *K* clusters, minimizing the sum of within-cluster distance. We varied the value of *K* between 2 and 20 and calculated the amount of across-subject variance explained in each case using the approach described in our previous publications^13–15^ and identified the optimal value of *K* when the variance saturates. We further confirmed that the percentage gain in variance in going from *K-1* to *K* clusters for each value of *K* beyond the optimal value remained below 0.5. Subsequently, for all m voxels, the z-scored BOLD signals were averaged across all time frames belonging to each cluster, resulting in *K* combined CAPs.

Next, we performed a one-sample two-tailed T-test to test the mean activation, across the occurrences of each CAP in the combined image-series of each voxel against a null hypothesis of zero activation and corrected for multiple comparisons using the Bonferroni correction (p<0.01). The voxels that showed significant activation or deactivation constituted the one-sample T-statistic maps for each CAP, representing the regional activities that prominently constitute the CAP. For each of the CAPs, we calculated the temporal and spatial properties:

1. Occurrence percentage: the percentage of the time frames corresponding to a CAP within a subject’s image-series.
2. Duration: number of consecutive frames corresponding to a CAP, averaged across all its occurrences, within a subject.
3. Voxel-level activations: Mean activation across all occurrences of a CAP within the image series for each of the four combinations of a genotypic group & age (4M WT, 4M TG, 10M WT, and 10M TG). We also calculated voxel-level activations in each subject by averaging across all occurrences of the CAP in each subject for the partial least squares regression analysis (See the section on relationship between BBMs, behavioral measures, and RS functional measures).

### Statistical analysis

For each behavioral measure, we assessed the main effects of genotype, time (acquisition day), and time x genotype interaction using a repeated measures two-way ANOVA. If no significant interaction effect was observed, it was removed from the model and the main effects were assessed. The post-hoc temporal comparisons were made between acquisition days 2 to 10 with the first day of acquisition.

Repeated measures two-way ANOVA tests were also used to assess the age, genotype, and age x genotype effect on ROI-level FC, network FC, as well as duration and occurrence percentage of CAPs. Effects of genotype, age, and genotype x age on spatial activation within each CAP were assessed for each voxel that showed a significant (de)activation in the combined 1-sample T-map using a two-way ANOVA test. If no significant interaction effect was observed, it was removed from the model and the main effects were assessed. All post-hoc comparisons were corrected for multiple comparisons using the Benjamini Hochberg method to control false discovery rate (FDR).

Data for BBMs Aβ38, Aβ40, and Aβ42 were assessed for normality using the Kolmogorov–Smirnov test. For Aβ38, data from wild-type and transgenic groups were normally distributed and compared using a paired t-test. As Aβ40 and Aβ42 data did not meet the assumption of normality in the WT group, group comparisons were performed using the non-parametric Mann–Whitney U test.

### Relationship between BBMs, behavioral measures and RS functional measures

We used partial least squares regression (PLSR) to investigate the relationship between behavioral measures, blood biomarkers and RS-FC, and spatial properties of RS-CAPs in the TG animals. Two behavioral measures - reference and working memory errors on the last day (day 10) of the acquisition period were considered as response/explained variables. BBMs included *Aβ*38, *Aβ*40, and *Aβ*42 values. ROI-based pairwise FC (561) values, within- and between-network FCs (15 in total), and voxel-wise activations for significantly (de)activated voxels for all six CAPs at 4 and 10 months were used as the predictors for each of the three behavioral measures as well as each BBM. Similarly, BBMs were used as predictors of behavioral measures. Values of each predictor and response variable were WT-normalized, thus for each TG subject, the mean of the (age-matched) WT group was subtracted from its value for each variable and divided by the standard deviation in the (age-matched) WT group.

PLSR is a multivariate regression technique that is well suited when there are more variables than observations (subjects in this case) and there could be high collinearity between the predictor variables. In the case of this manuscript, both these conditions apply, especially when RS-functional measures were used as predictor variables (whether FC values or CAPs’ spatial properties). It identifies components from both the predictor and response variables so that the covariance between the predictor and response variables is maximally accounted for. We used leave-one-out cross-validation to obtain the optimal number of components for each fitted model and calculated the most variance (𝑅^2^) explained by the following expression:

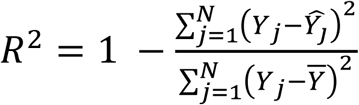

Here, N is the number of subjects; 𝑌_𝑗_ is the value of the response variable for subject *j*, *Ŷ*_*j*_ is the value predicted by the model for subject *j* and *Ȳ* is the mean value of the response variable across all subjects. We also calculated the 95% confidence intervals for the 𝑅𝑅^2^ values from 1000 bootstraps (with replacement) and after correcting for the number of predictor variables by controlling for the family-wise-error rate (FWER). Fitted models also yielded the regression coefficients, *β* values, and their 95% confidence intervals corrected for the number of predictor variables by controlling FWER. The variables for which the confidence intervals for *β* values did not cross zero were considered salient. TG animals with high/low values of the response variables (amyloid markers, reference and working memory errors) we considered in this study correspond to high/low levels of severity of the phenotype vis-à-vis the WT. However, high values of FC and CAP activations in TG animals do not necessarily imply that functional change in these parameters is more severe than low values of FC and/or CAP activations. Therefore, to ensure the interpretation that positive and negative *β* weights imply higher values of response variables correspond to greater and lower severity, respectively, in the functional variables, we flipped the signs of weights for measures for which the mean value in the TG animals was lower than in the WT animals. Then we plotted, in case of ROI-FC, the top 20% of positive and negative *β* values in the matrix format to identify the salient predictors. In case of CAPs, we applied a 20-voxel cluster threshold to identify clusters of salient voxels in each slice.

## Author Contributions

**AAA:** Investigation, Formal analysis, Writing – Reviewing and Editing; **LB:** Investigation, Formal analysis, Writing - Reviewing and Editing; **SDW:** Investigation, Data Acquisition, Writing - Reviewing and Editing; **SVH:** Investigation; **JG:** Methodology, Software; **IDW:** Methodology, Writing - Reviewing and Editing; **MV:** Conceptualization, Supervision, Funding Acquisition, Project Administration, Writing – Reviewing and Editing; **MHA:** Conceptualization, Supervision, Funding Acquisition, Methodology, Formal Analysis, Writing – Original Draft, Reviewing and Editing.

## Funding

This study was supported by the Fund of Scientific Research Flanders (FWO-G045420N) and the Stichting Alzheimer Onderzoek (SAO - FRA 2020/027) awarded to MV, and BOF – small research grant (KP-BOF 2024 / ID: 50751) awarded to MHA. The computational resources and services used in this work were provided by the HPC core facility CalcUA of the University of Antwerp, the VSC (Flemish Supercomputer Center), funded by the Hercules Foundation, and the Flemish Government department EWI. The work was also supported by the Flemish Impulse funding for heavy scientific equipment under grant agreement number OZ3544 (granted to Annemie Van der Linden).

## Data Availability statement

The datasets used and/or analyzed during the current study will be made publicly available within six months from the publication of the paper. In the meantime, they can be obtained by submitting a request to the corresponding author.

## Supplementary Figures

**Supplementary Figure 1:**
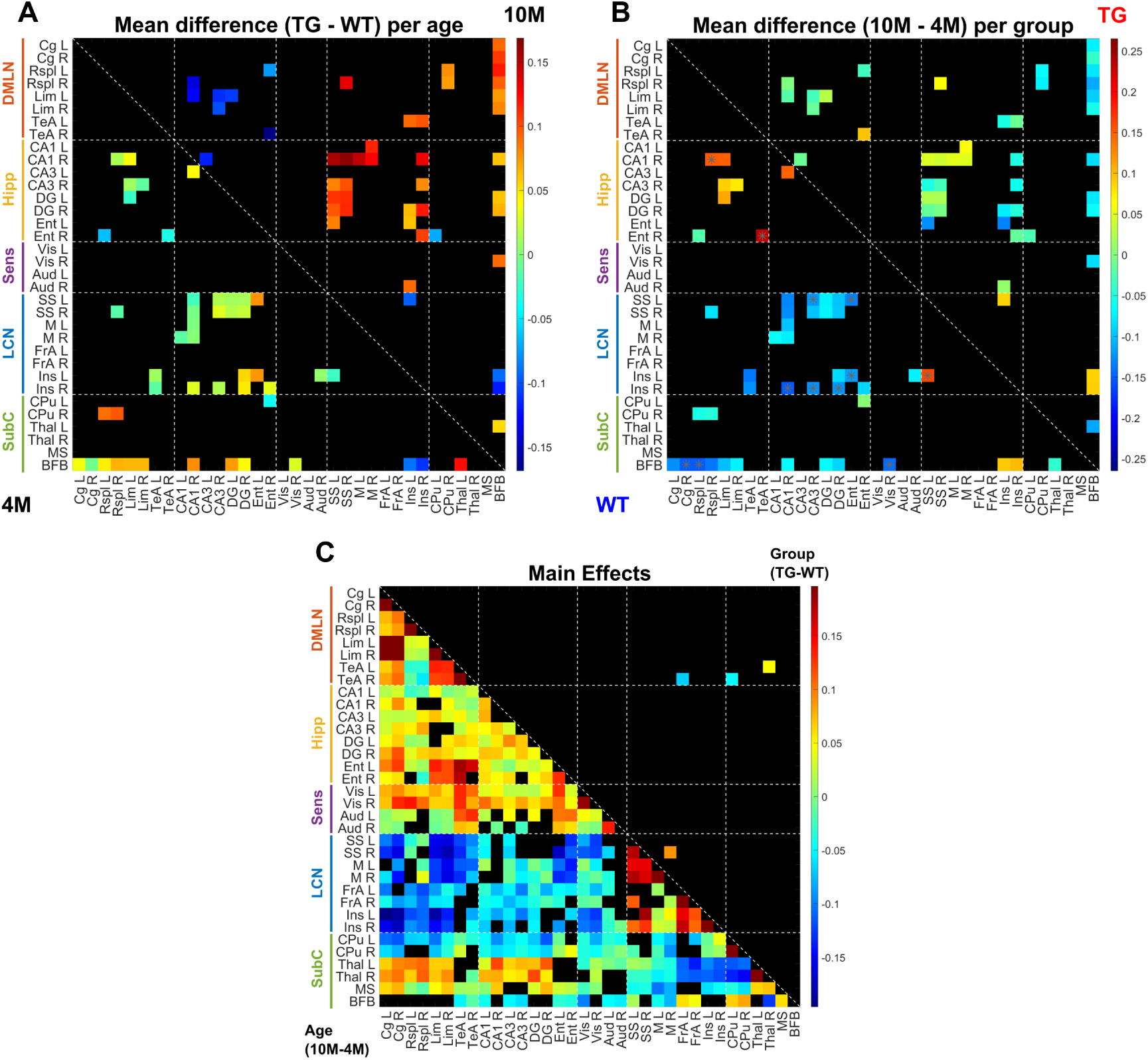
**A, B:** Inter-genotype difference (A; 4M/10M: below/above the diagonal) and inter-age difference (B; WT/TG: below/above the diagonal) in FC for ROI-pairs that showed a significant genotype*age interaction effect (p < 0.05, repeated measures two-way ANOVA). **C**: Inter-age difference (below the diagonal) and inter-genotypic group difference (above the diagonal) in FC for ROI-pairs that showed a significant (p < 0.05, repeated measures two-way ANOVA) main effect of age and genotype respectively. Asterisk: p < 0.05, FDR corrected for all ROI pairs found to be significant for each effect.

**Supplementary Figure 2:**
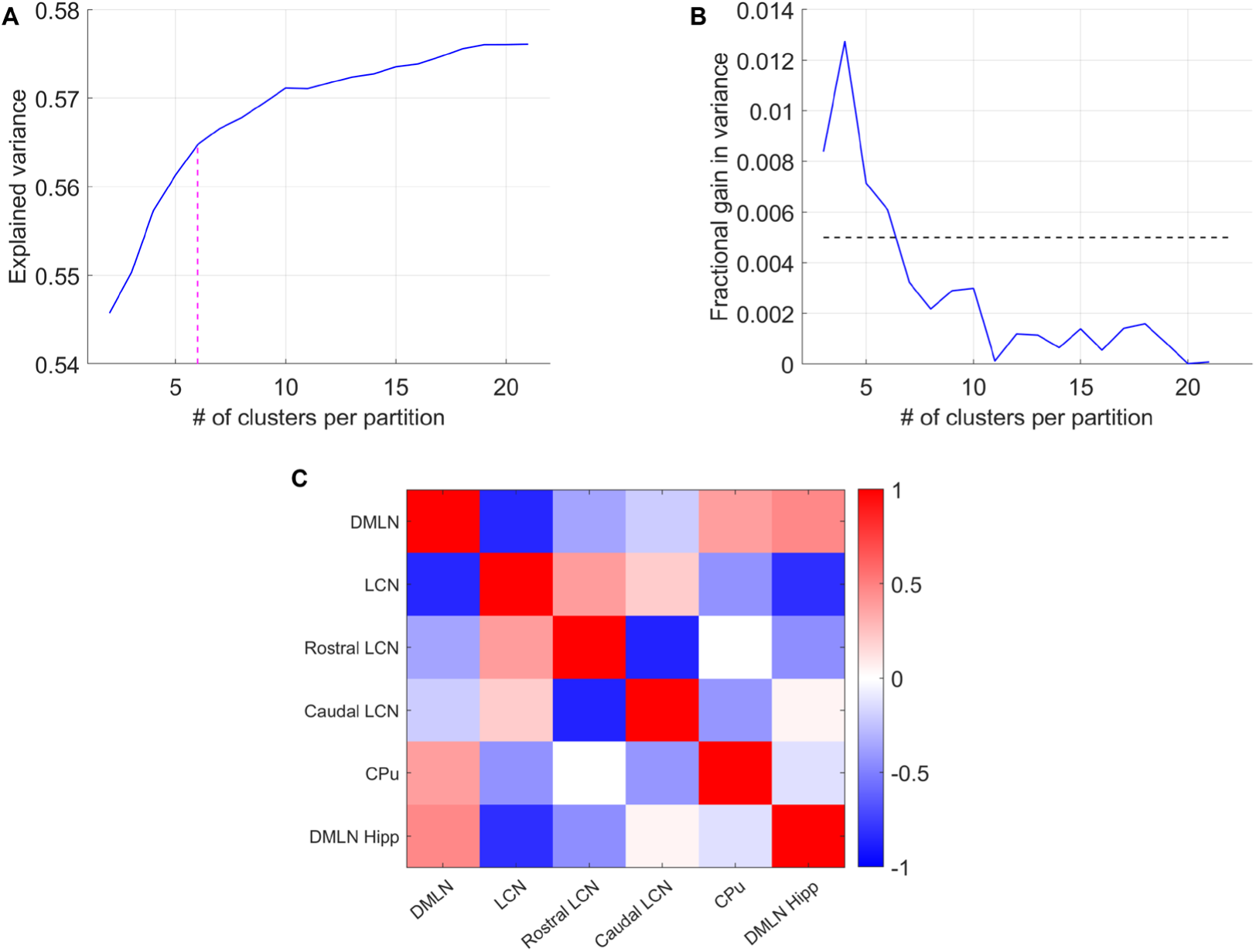
**A:** Variance in the concatenated image-series explained by different numbers of clusters. Magenta vertical line shows the optimal number of clusters corresponding to the elbow point of this curve. **B:** Fractional gain in variance in going from k-1 to k clusters as a function of number of clusters. **C:** Pearson’s correlation between spatial patterns of every pair of CAPs shows the first four CAPs forming two anticorrelated pairs while the last two CAPs do not show high anticorrelation.

**Supplementary Figure 3:**
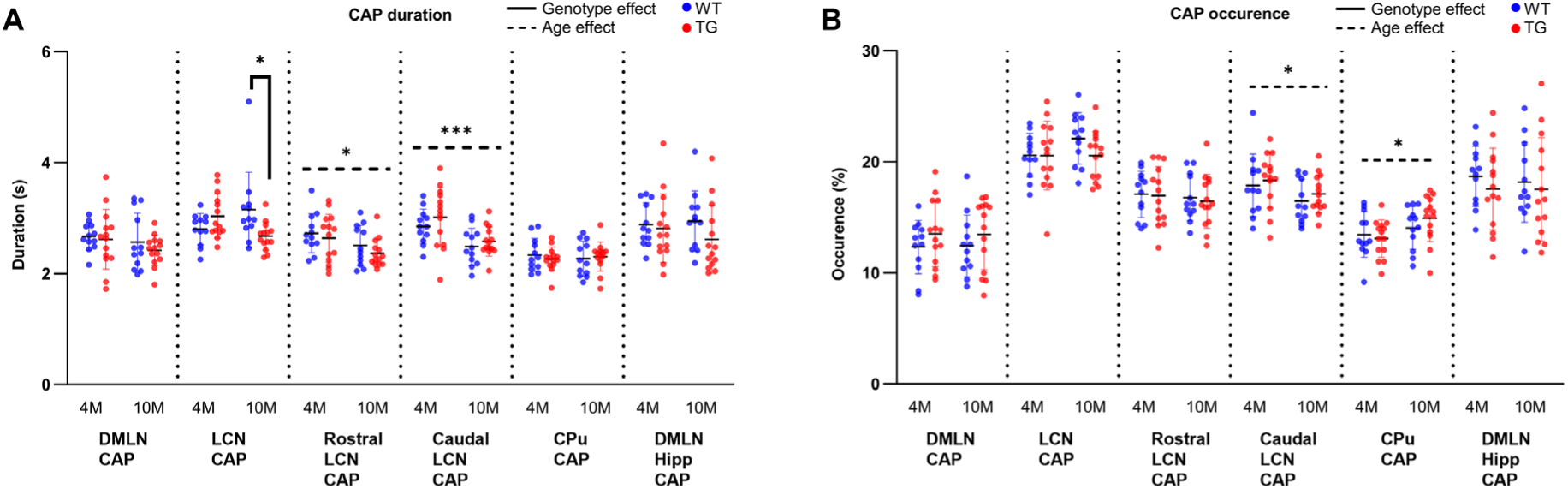
Age, genotypic group, and interaction effects on CAP duration (A), and occurrence (B) using repeated measures two-way ANOVA followed by appropriate post-hoc comparisons. *: p < 0.05; ***: p < 0.001.

**Supplementary Figure 4:**
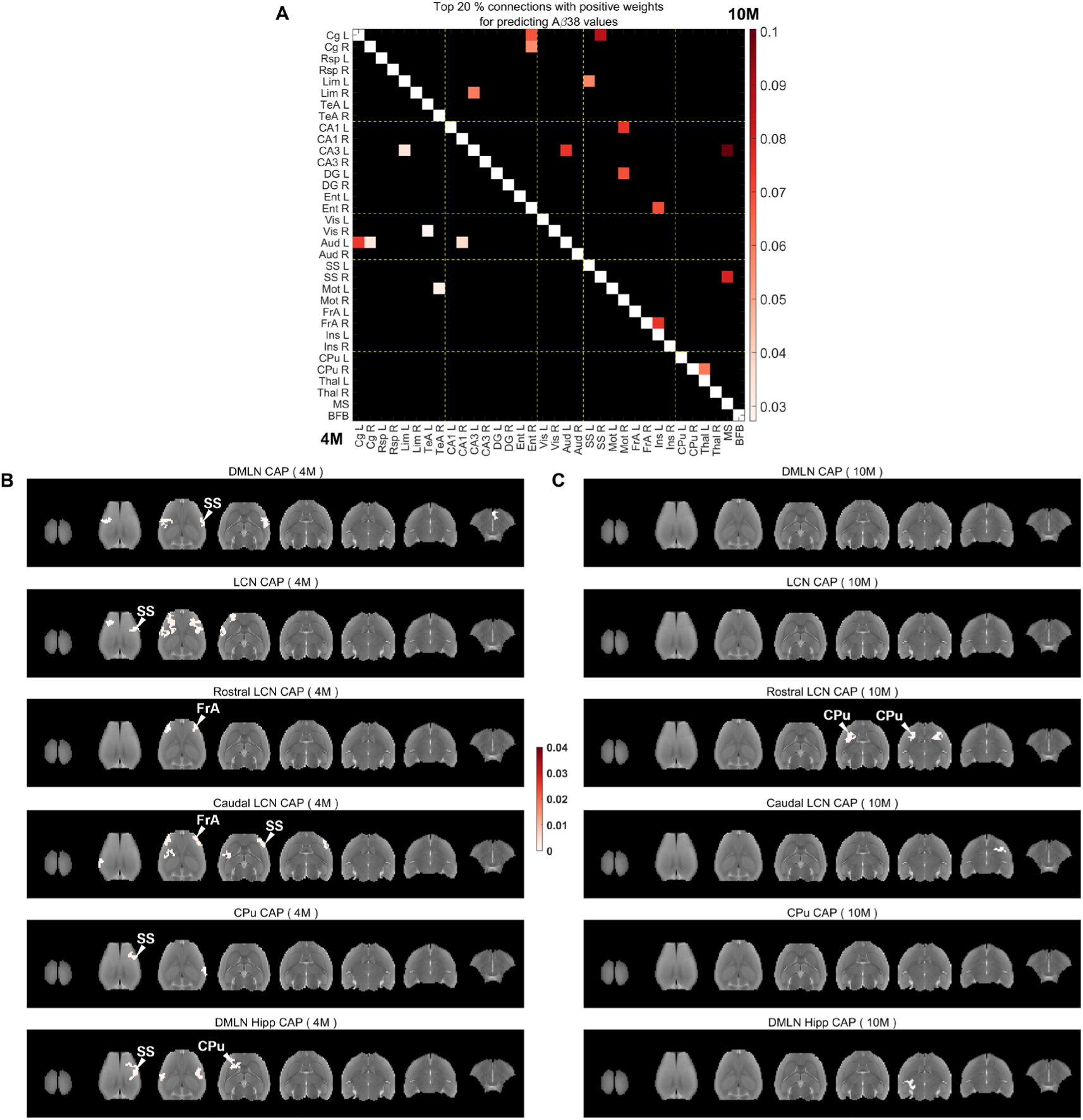
Salient functional connections (A) and voxel-clusters (B-C) within CAPs with positive beta weights at pre-plaque and plaque stages that explain *Aβ*38 values in the TG animals. **A:** Top 20% highest positive beta weights of functional connections with significant 99% confidence intervals at 4M (below the diagonal) and 10M (above the diagonal). **B, C:** Positive beta weights for salient clustered voxels (cluster-size: 20 voxels, in plane), for which FWER corrected 95% confidence intervals do not cross zero, in each CAP at 4M (B) and 10M (C).

**Supplementary Figure 5:**
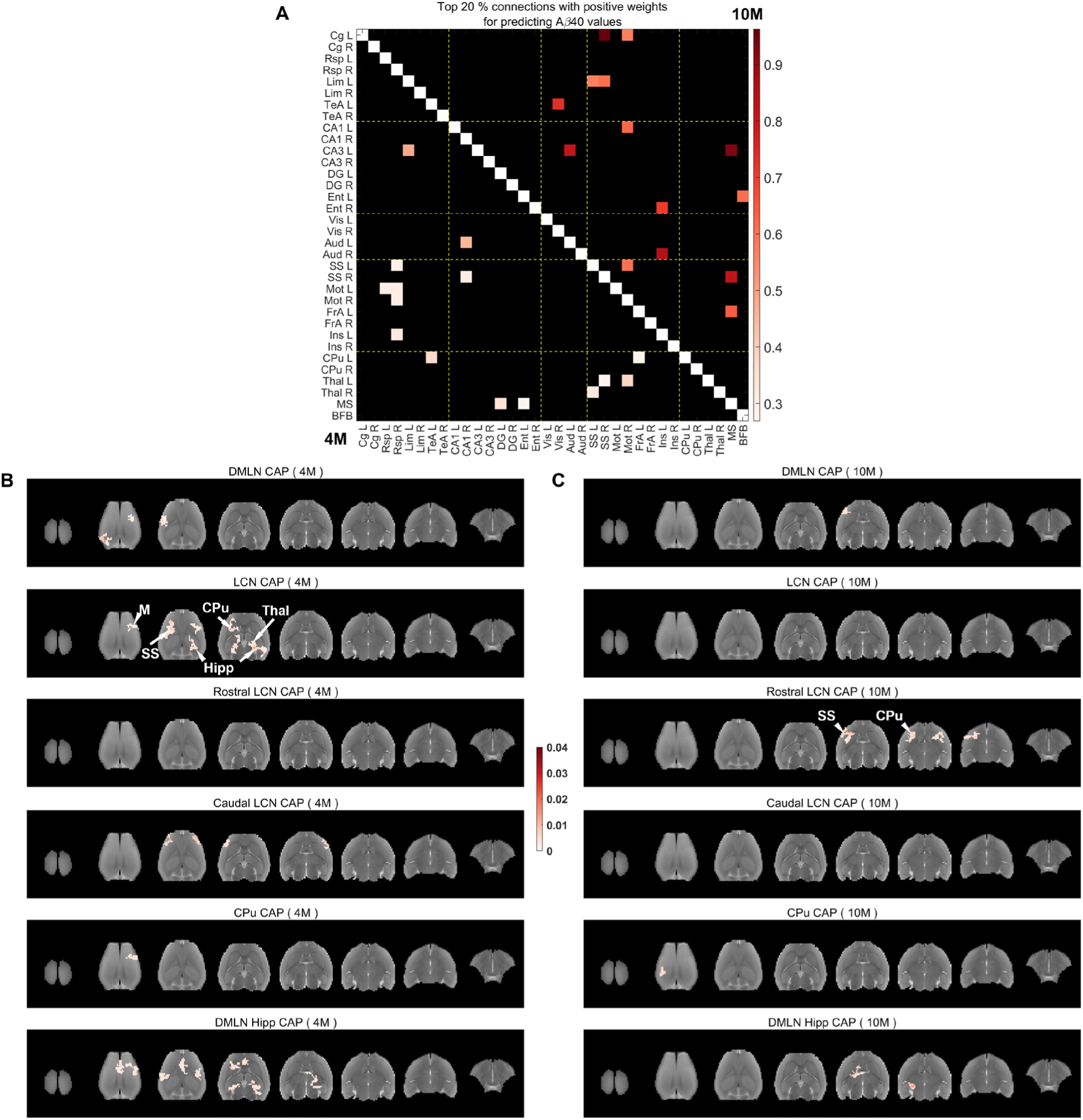
Salient functional connections (A) and voxel-clusters (B-C) within CAPs with positive beta weights at pre-plaque and plaque stages that explain *Aβ*40 values in the TG animals. **A:** Top 20% highest positive beta weights of functional connections with significant 99% confidence intervals at 4M (below the diagonal) and 10M (above the diagonal). **B, C:** Positive beta weights for salient clustered voxels (cluster-size: 20 voxels, in plane), for which FWER corrected 95% confidence intervals do not cross zero, in each CAP at 4M (B) and 10M (C).

**Supplementary Figure 6:**
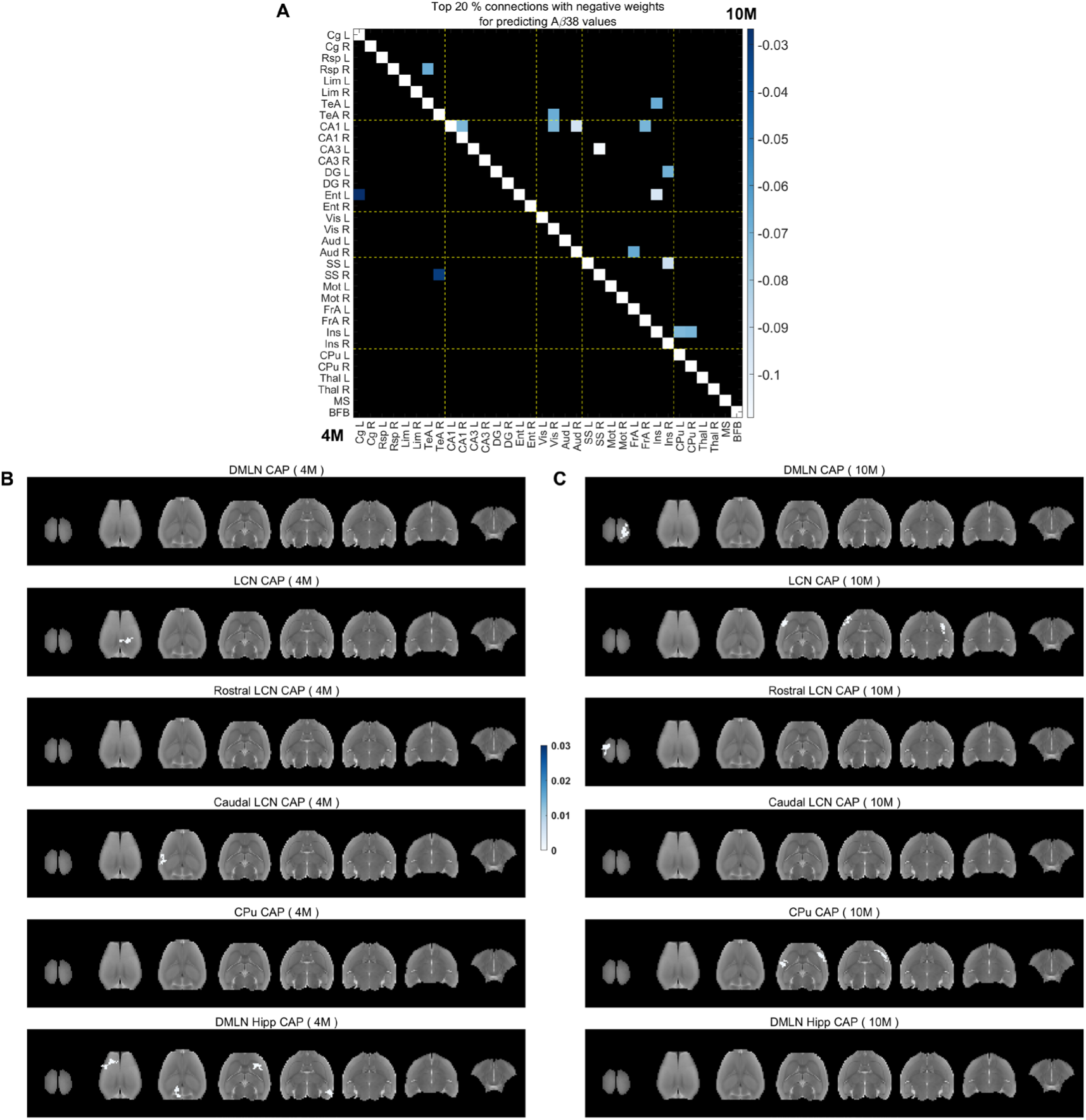
Salient functional connections (A) and voxel-clusters (B-C) within CAPs with negative beta weights at pre-plaque and plaque stages that explain *Aβ*38 values in the TG animals. **A:** Top 20% highest negative beta weights of functional connections with significant 99% confidence intervals at 4M (below the diagonal) and 10M (above the diagonal). **B, C:** Negative beta weights for salient clustered voxels (cluster-size: 20 voxels, in plane), for which FWER corrected 95% confidence intervals do not cross zero, in each CAP at 4M (B) and 10M (C).

**Supplementary Figure 7:**
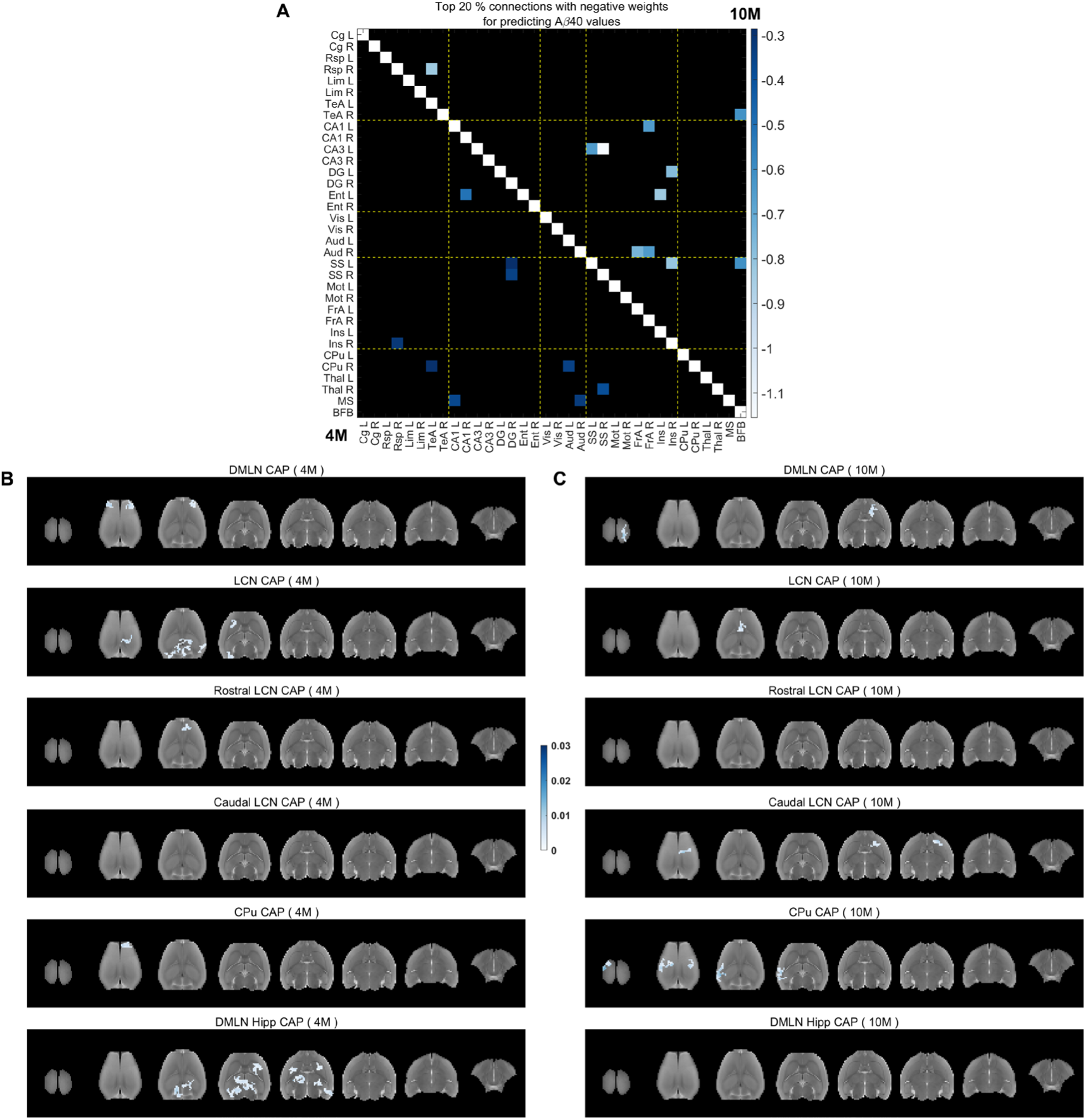
Salient functional connections (A) and voxel-clusters (B-C) within CAPs with negative beta weights at pre-plaque and plaque stages that explain *Aβ*40 values in the TG animals. **A:** Top 20% highest negative beta weights of functional connections with significant 99% confidence intervals at 4M (below the diagonal) and 10M (above the diagonal). **B, C:** Negative beta weights for salient clustered voxels (cluster-size: 20 voxels, in plane), for which FWER corrected 95% confidence intervals do not cross zero, in each CAP at 4M (B) and 10M (C).

**Supplementary Figure 8:**
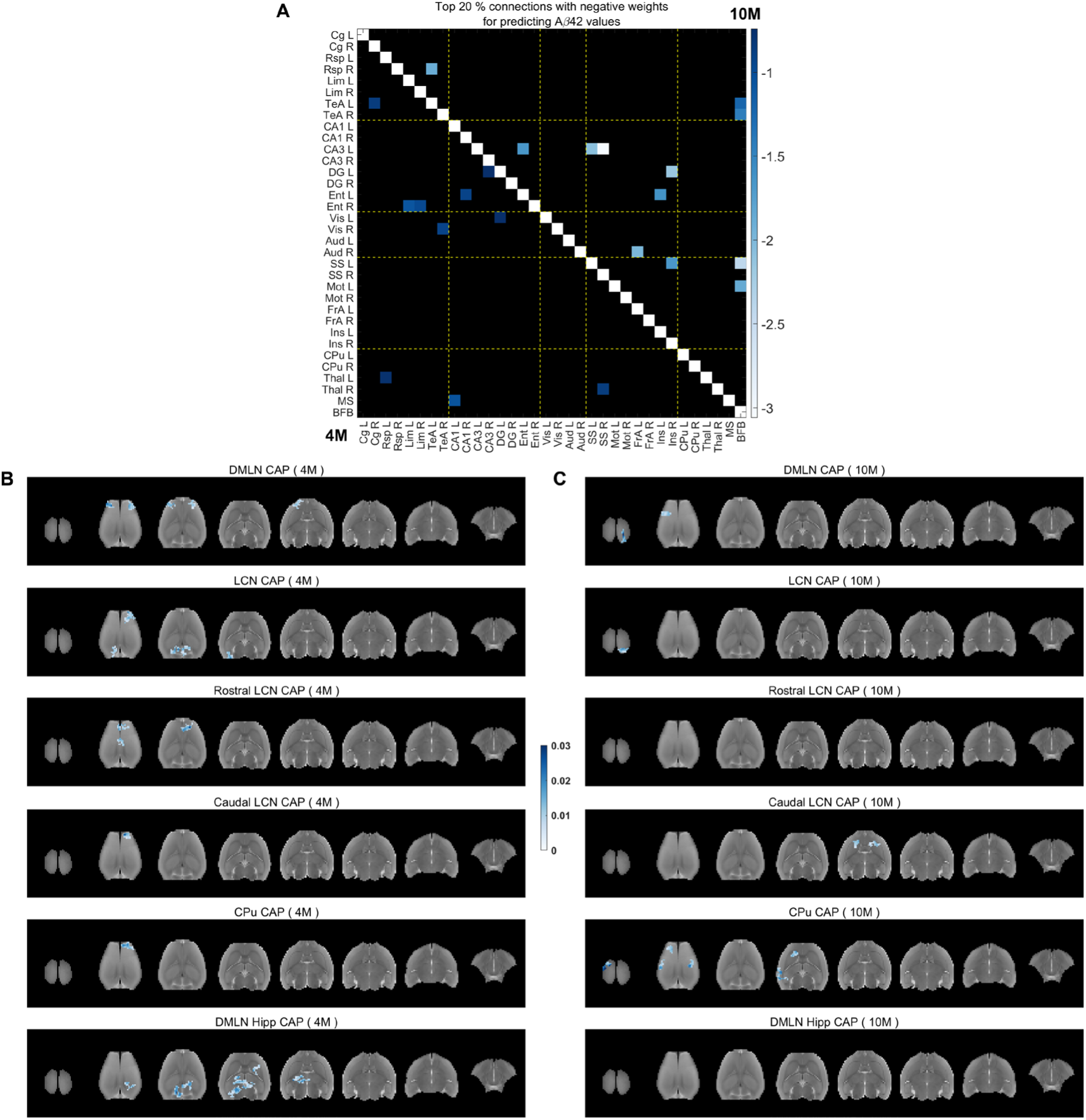
Salient functional connections (A) and voxel-clusters (B-C) within CAPs with negative beta weights at pre-plaque and plaque stages that explain *Aβ*42 values in the TG animals. **A:** Top 20% highest negative beta weights of functional connections with significant 99% confidence intervals at 4M (below the diagonal) and 10M (above the diagonal). **B, C:** Negative beta weights for salient clustered voxels (cluster-size: 20 voxels, in plane), for which FWER corrected 95% confidence intervals do not cross zero, in each CAP at 4M (B) and 10M (C).

**Supplementary Figure 9:**
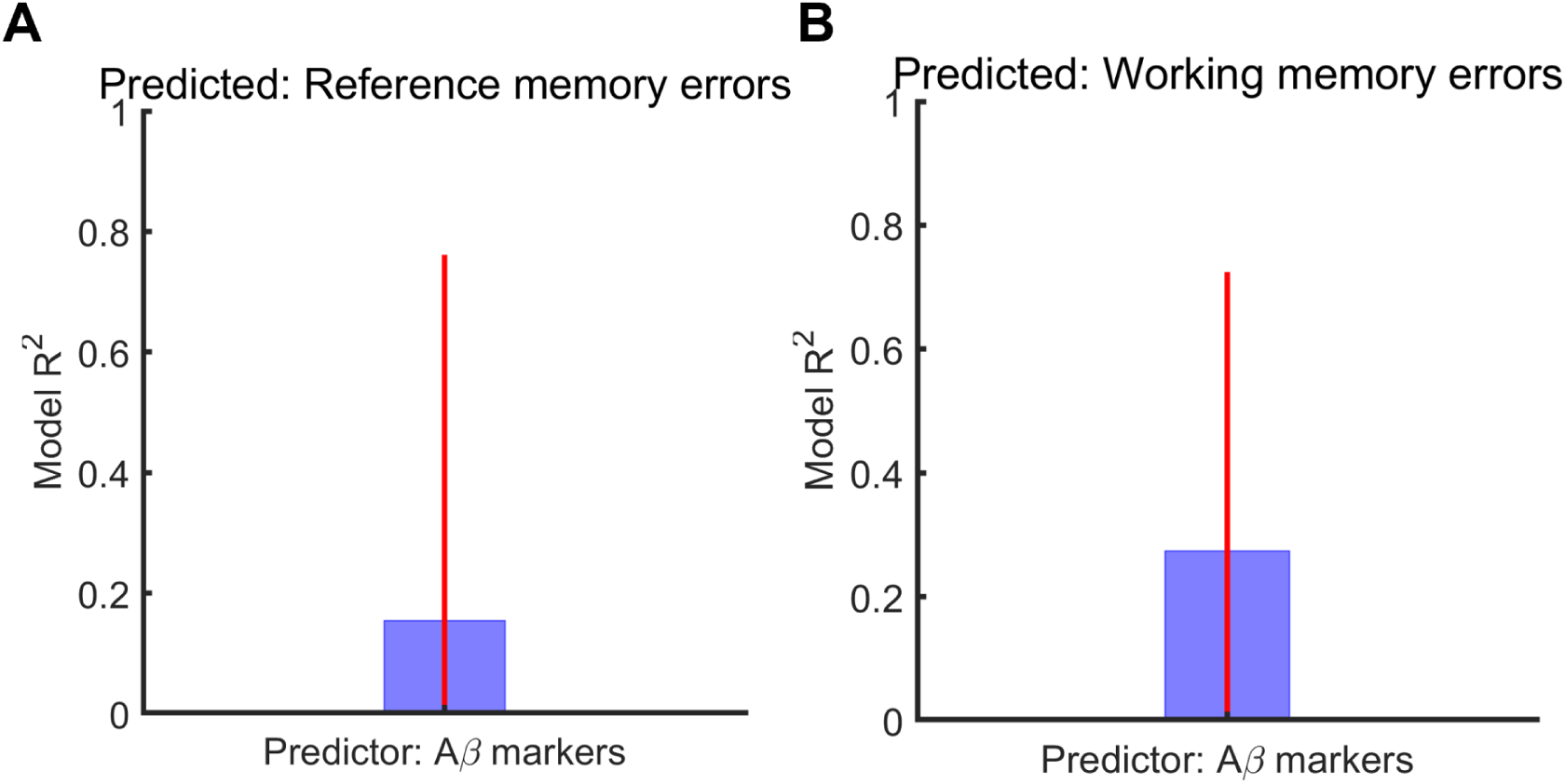
PLSR model *R*^2^ for predicting the performance on memory tasks (reference and working memory errors) on acquisition day 10 measured during the plaque stage (10.5M) using plasma levels of *Aβ*38, *Aβ*40, *Aβ*42 at the plaque (11M) stage. Here we used WT-normalized values for each amyloid marker to explain the WT-normalized values of reference memory errors (A), and working memory errors (B) in each individual TG subject respectively. Red vertical lines: family-wise error (FWE) - corrected 95% confidence intervals.

**Supplementary Figure 10:**
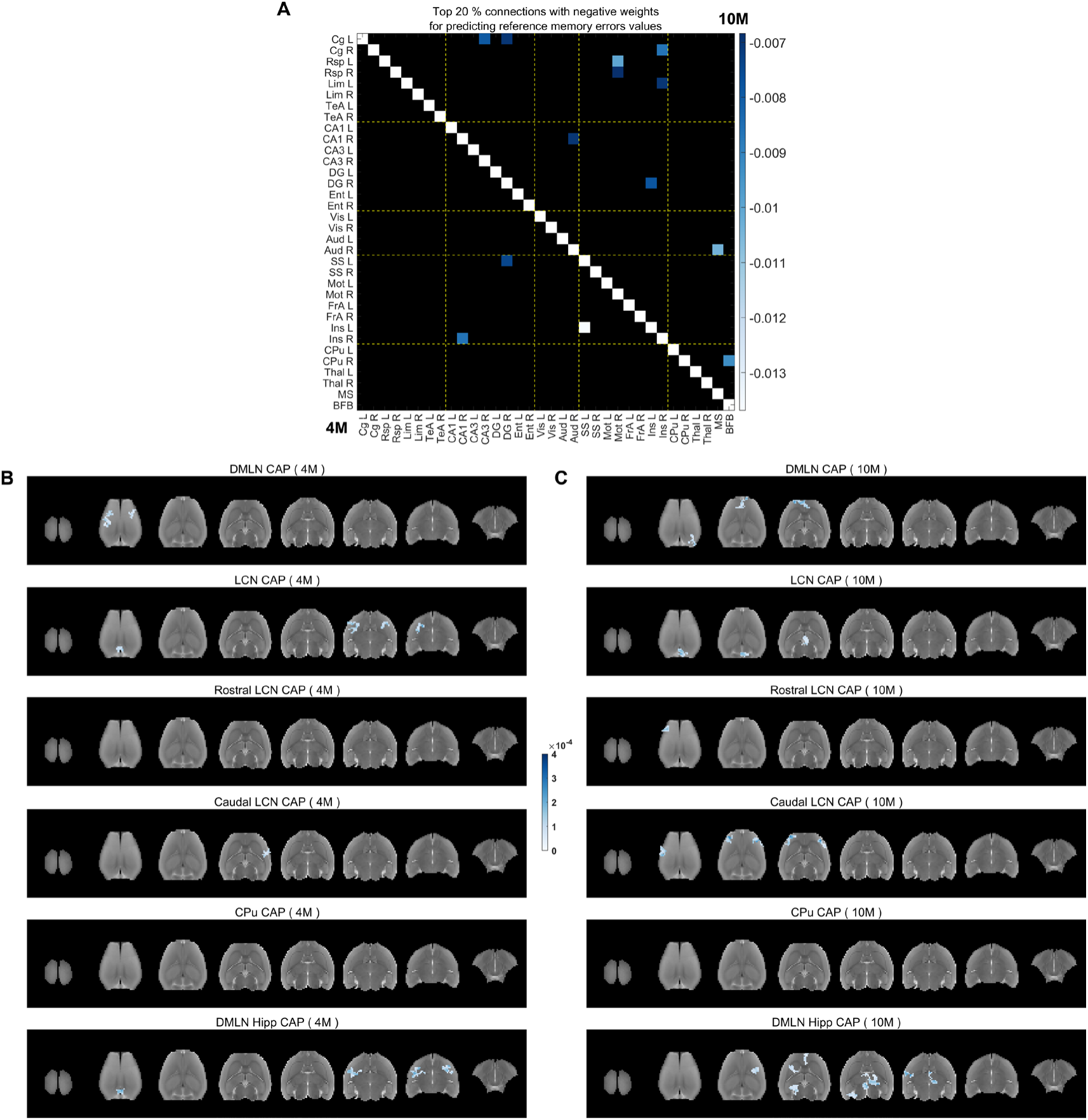
Salient functional connections (A) and voxel-clusters (B-C) within CAPs with negative beta weights at pre-plaque and plaque stages that explain WT-normalized reference memory errors in the TG animals. A: Top 20% highest negative beta weights of functional connections with significant 99% confidence intervals at 4M (below the diagonal) and 10M (above the diagonal). **B, C:** Negative beta weights for salient clustered voxels (cluster-size: 20 voxels, in plane), for which FWER corrected 95% confidence intervals do not cross zero, in each CAP at 4M (B) and 10M (C).

**Supplementary Figure 11:**
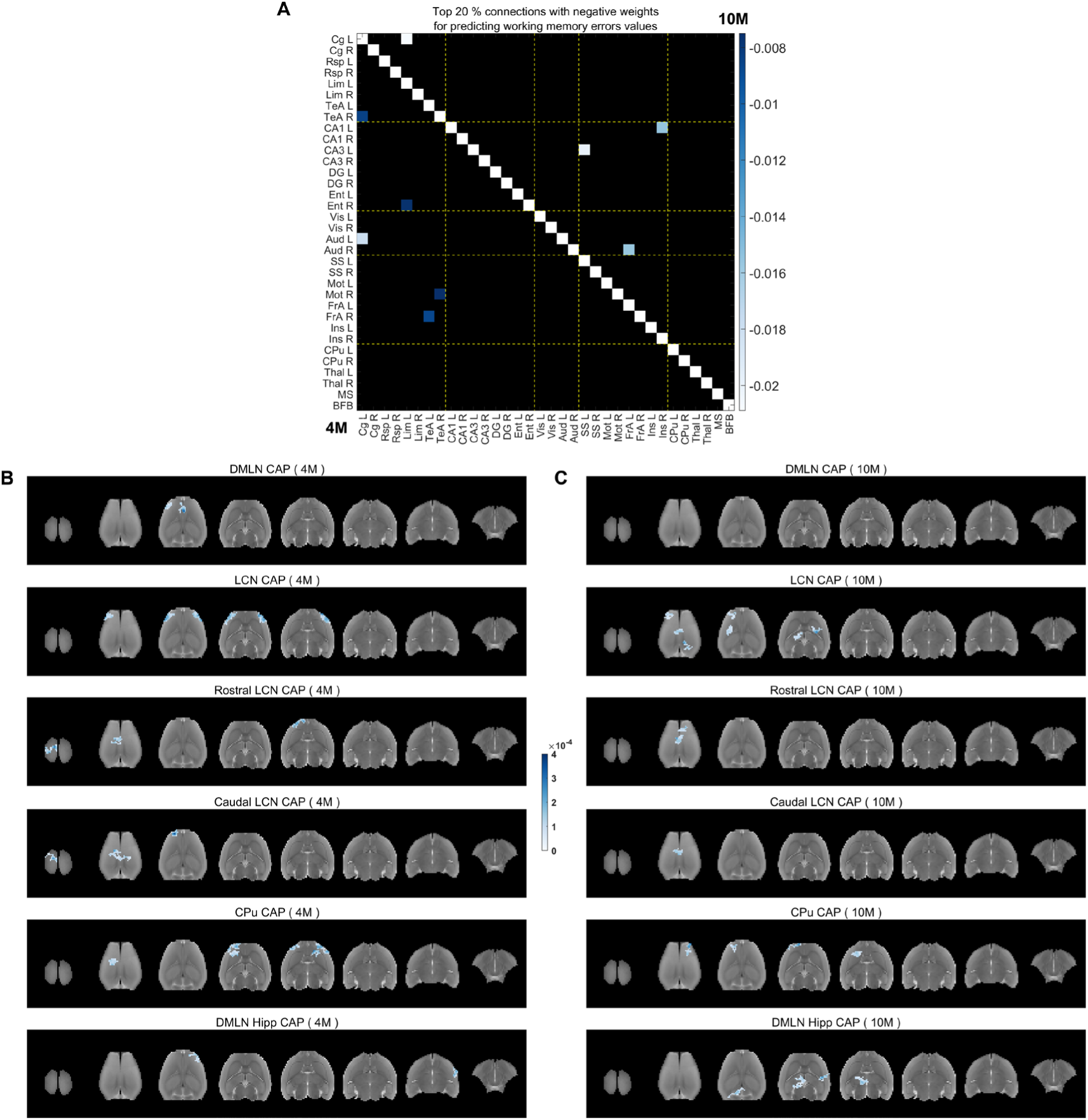
Salient functional connections (A) and voxel-clusters (B-C) within CAPs with negative beta weights at pre-plaque and plaque stages that predict WT-normalized working memory errors in the TG animals. A: Top 20% highest negative beta weights of functional connections with significant 99% confidence intervals at 4M (below the diagonal) and 10M (above the diagonal). **B, C:** Negative beta weights for salient clustered voxels (cluster-size: 20 voxels, in plane), for which FWER corrected 95% confidence intervals do not cross zero, in each CAP at 4M (B) and 10M (C).

## Notes

### Competing Interest Statement

The authors have declared no competing interest.

